# Intense storms affect sinking particle fluxes after the North Atlantic diatom spring bloom

**DOI:** 10.1101/2024.01.11.575202

**Authors:** Elisa Romanelli, Sarah Lou Carolin Giering, Margaret Estapa, David A. Siegel, Uta Passow

**Author notes:** **Corresponding author:** Elisa Romanelli, Interdepartmental Graduate Program in Marine Science University of California, Santa Barbara, CA.

## Abstract

The sinking of large particles (i.e., marine snow) has long been recognized as a key pathway for efficient particulate organic carbon (POC) export to the ocean interior during the decline of spring diatom blooms. Recent work has suggested that particles smaller than marine snow can also substantially contribute to POC export. However, a detailed characterization of small and large sinking particles at the end of blooms is missing. Here, we separately collected suspended and small and large sinking particles using Marine Snow Catchers and assessed their biogeochemical composition after the North Atlantic spring bloom in May 2021. During the three weeks of sampling, when four intense storms (maximum wind speeds 37 – 50 kts) created high turbulent energy dissipation rates and deepened the mixed layer, we observed two distinct sedimentation episodes. During the storm periods, sinking particles were dominated by small (diameter < 0.1 mm), slow-sinking (~18 m d^−1^), silica-rich particles that carried a moderate POC flux (< 6 mmol C m^−2^ d^−1^) to 500 m depth. Once the storms ceased, the volume of large (diameter > 0.1 mm), fast-sinking (> 75 m d^−1^), carbon-rich marine snow aggregates (not fecal pellets) increased exponentially and POC fluxes at 100 m depth were more than fourfold greater (30±12 mmol C m^−2^ d^−1^) than those during the previous event. The aggregates consisted of a mixed post-bloom plankton community. Our data suggest that the intense storms determined the timing, type, and magnitude of POC flux at the end of a spring phytoplankton bloom.

## Introduction

Spring phytoplankton blooms account for a significant fraction of the annual oceanic primary production and substantially contribute to carbon sequestration by the ocean (Henson et al., 2019; Siegel et al. 2023). The termination of a diatom bloom is frequently viewed as a single event of rapid diatom aggregation, triggered when nutrient limitation causes bloom senescence, and subsequent sedimentation of carbon-rich, fast-sinking diatom aggregates (sinking velocity > 50 m d^−1^) (Alldredge and Silver 1988; Alldredge and Gotschalk, 1989; Riebesell, 1991b; Kiørboe et al., 1996). The result is the efficient export of particulate organic carbon (POC) to the ocean interior (Honjo, 1982; Smetacek, 1985). The transition between bloom growth and export via marine snow is difficult to predict and observe in the open ocean due to its ephemeral and spatially heterogeneous nature. We therefore lack a detailed understanding of the spatiotemporal dynamics of sinking particles during the declining phase of a diatom bloom.

Marine snow aggregates associated with diatom bloom termination form via coagulation of particles in the presence of high particle concentrations (e.g., phytoplankton) and increased levels of stickiness that enable attachment of particles that collide (Jackson, 1990). Marine snow aggregates are composed of microbial cells and organic and inorganic particulate material embedded in a matrix of transparent exopolymer particles (TEP). Transparent exopolymer particles are gel-like particles largely composed of polysaccharides exuded as extracellular, surface-active exopolymers (Passow, 2002), especially under nutrient-limited conditions (Obernosterer and Herndl, 1995). Due to their sticky nature, the presence of TEP favors the formation of aggregates by increasing stickiness, that is the probability that the collision of particles will result in coagulation (Passow and Alldredge, 1994; Jackson, 1995). However, TEP are inherently positively buoyant (Engel and Schartau, 1999; Azetsu-Scott and Passow, 2004), and their role in regulating the magnitude of POC export is variable and dependent on the oceanic ecosystem (Mari et al., 2017; Romanelli et al., 2023). Another important factor controlling the formation of aggregates is the intensity of turbulent shear, which needs to be large enough to promote particle-particle collisions (Jackson, 1990; Stemmann et al., 2004). In the open ocean, turbulent kinetic energy dissipation rates (*ϵ*) in the mixed layer vary between 10^−10^ W kg^−1^ to 10^−1^ W kg^−1^ (D’Asaro, 2014), with the highest rates occurring under intense storm conditions near the sea surface. However, when turbulent dissipation rates are large (*ϵ* > 10^−6^ W kg^−1^), disaggregation of marine snow may be more important than aggregation in dictating marine snow size (Dyer and Manning, 1999; Ruiz and Izquierdo, 1996; Takeuchi et al., 2019). The impact of storms on sedimentation of marine snow aggregates has rarely been observed in situ (D. A. Siegel et al., unpubl.).

Here we present detailed observations of marine particle dynamics immediately after the main sedimentation event of a spring diatom bloom. This period was characterized by a series of intense storms that impacted sinking particle fluxes. To understand particle sedimentation under this post-bloom scenario, we characterized suspended, small sinking particles and fast-sinking marine snow collected using Marine Snow Catchers (MSC; Lampitt et al., 1993) during the NASA-led Export Processes in the Ocean from RemoTe Sensing (EXPORTS) North Atlantic field campaign, which provided a broad suite of contextualizing observations (Johnson et al., 2023).

## Methods

### Study site

Observations of suspended and sinking particle characteristics were obtained in spring 2021 (May 4 to 29, 2021) on board the RRS James Cook, which targeted biogeochemical and ecological processes within an anticyclonic eddy (Johnson et al., 2023). The goal of sampling within a particle retentive eddy was to obtain time series measurements in a Lagrangian manner. The eddy was located ~170 km east of the site of the Porcupine Abyssal Plain Sustained Observatory (PAP-SO) (Erickson et al. 2023). During the field campaign four windstorms substantially deepened the mixed layer (ML, Δ*ρ* threshold of +0.05 kg m^−3^) by 25–40 m (**Figure 1**) and exchanged surface water masses with the surrounding waters due to Ekman transport (Johnson et al., 2023). For this reason, the eddy was deemed retentive only below a depth of ~100 m (eddy core waters) and within 15 km from its center (Johnson et al., 2023). The deployments of the MSCs were performed at 7 ± 4 km from the eddy center except on May 12 where samples were collected at 17 km. To better investigate temporal trends, we grouped our observations in five time periods (P) demarcated by the four storms (P1: May 4–7; P2: May 12– 14; P3: May 16–19; P4: May 22; P5: May 25–29; **Figure 1**). The first storm (May 7–10) exchanged approximately 73 % of the surface waters above the eddy with surrounding waters, and we hence refrain from drawing conclusions regarding temporal trends between P1 and P2 above the eddy core waters, but P1 observations are still included in the results and figures. The estimated exchange above the ML caused by the three subsequent storms was less than 50 % for each (Johnson et al., 2023); hence, we will interpret P2 to P5 as a time series.

**Figure 1.**
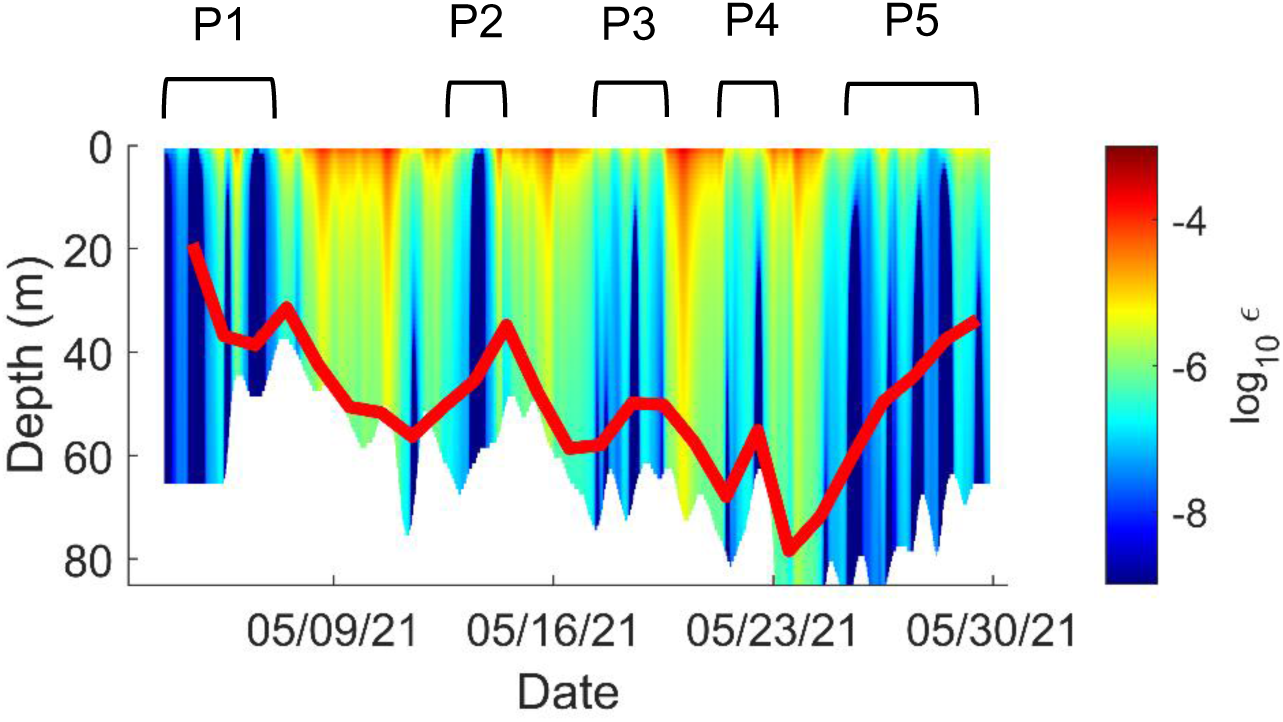
Estimates of the turbulent kinetic energy dissipation rate (*ϵ*, W kg^−1^) within the near-surface mixed layer during the field campaign. Red line represents the depth of the daily mean mixed layer. The five sampling periods (P) were demarcated by the storms (P1: May 4–7; P2: May 12–14; P3: May 16–19; P4: May 22; P5: May 25–29).

### Collection of suspended and sinking particles

Suspended and sinking particles were collected at 16 stations with four 100-liter Marine Snow Catchers (MSC, OSIL), at depths between 40 m and 500 m. All samples were collected in the morning (03:00–10:00 UTC, local time) except for two casts (on May 13 and May 25) collected at 16:30 UTC. At each station, one MSC was deployed 10 m below the ML, one 110 m below the ML, one at 300 m or 500 m and one at a variable depth matching the deployment depths of the Neutrally Buoyant Sediment Traps (NBST, Estapa et al., 2023).

The MSC methodology employed here is described in detail by Romanelli et al. (2023). Briefly, the MSC is a water sampler with a top and base section (82.2 L and 7.6 L, respectively) that allows the collection of particles according to their sinking velocity (Riley et al., 2012; Giering et al., 2016). To separately collect slow- and fast-sinking particles, a circular plastic tray (polypropylene, diameter: 18.5 cm; height: 4 cm; volume: ~1 L) was placed at the bottom of the MSC’s base. We defined three particle fractions according to their sampling location within the MSC following 2 hours of settling: *top* (*t*), *base* (*b*) and *tray* (*tr*). The *top* fraction was collected from the central tap and contained suspended particles. After draining and removing the top section, water overlying the tray in the base section was collected constituting the *base* particle fraction. The particle fraction in the tray was designated as the *tray* fraction. At 9 stations, fast-sinking marine snow particles (defined here as ESD >0.1 mm, examples in Supporting Information **Fig. S1**) were removed from the tray via manual pipetting, visually classified with a ruler (± 10 %) into size bins of 0.5–1, 1–2, 2–4, and 4–8 mm (ESD) and preserved in 4 % formaldehyde to perform microscopy later (**Table 1**). On May 27, fast-sinking marine snow was removed from the tray to measure its POC mass (**Table 1**) At 3 stations the entire particulate pool in the tray was processed as the *tray* fraction (i.e., marine snow was not removed) to enable estimates of the average sinking velocity of fast-sinking particles using the method detailed below (**Table 1**).

**Table 1.** Analyses performed at each station sampled. The last column indicates in which station marine snow was sized, counted, and removed from the MSC tray. On May 16 marine snow sample was lost. On May 27, marine snow was filtered to measure the POC and PON mass but not counted quantitatively in the tray.

| Date | Particulate organic carbon and nitrogen (POC and PON) | Total Particulate Carbon (TC) | Biogenic silica (bSi) | Lithogenic silica (lSi) | Sinking velocity of fast-sinking particles | Marine snow removed from the tray |
| --- | --- | --- | --- | --- | --- | --- |
| May 7 | × | × | × | × |  | × |
| May 12 | × | × | × |  |  | × |
| May 13 | × | × | × |  | × | × |
| May 14 | × |  |  |  |  |  |
| May 16 | × | × | × | × |  | × (No counts) |
| May 17 | × | × | × |  | × | × |
| May 18 | × |  |  |  |  |  |
| May 19 | × | × | × |  |  | × |
| May 22 | × | × | × |  |  | × |
| May 25 | × |  |  |  | × |  |
| May 26 | × | × | × |  |  | × |
| May 27 |  |  | × |  |  | × (Measured POC of marine snow) |
| May 28 | × | × | × | × |  | × |
| May 29 | × | × | × |  |  | × |

After removal of marine snow as appropriate (**Table 1**), the *top*, *base* and *tray* fractions were homogenized through shaking and subsampled using a graduated cylinder. Subsamples for POC, particulate organic nitrogen (PON), total particulate carbon (TC) and biogenic silica (bSi) were collected as described in section 2.4 (**Table 1**). In addition, the concentrations of POC and TEP in the mixed layer (21±5 m) were assessed at 7 stations located in the eddy core on samples collected with Niskin bottles mounted on a CTD-rosette. Analytical methods are detailed below. Table 1. Analyses performed at each station sampled. The last column indicates in which station marine snow was sized, counted, and removed from the MSC tray. On May 16 marine snow sample was lost. On May 27, marine snow was filtered to measure the POC and PON mass but not counted quantitatively in the tray.

### Biochemical analysis

Concentrations of POC and PON were determined by filtering duplicate subsamples onto pre-combusted (450 °C, 30 minutes) replicate GF/F filters (25 mm, Whatmann, UK). We filtered 1 L of the samples collected with the Niskin bottles, 1L of the MSC *top* fractions, 0.6–1 L of the MSC *base* fraction and 0.2 L of the MSC *tray* fraction. The filters were dried at 40°C, stored at room temperature and analyzed using a CEC 44OHA elemental analyzer (Control Equipment, US) after treatment with 150 μL of 10% HCl (v/v) to remove particulate inorganic carbon (PIC). Before calculating the concentrations of POC and PON, their measured masses were blank-corrected to account for contamination using the cruise-wide EXPORTS’ average (7.905 μg for POC; 1.782 μg for PON; n = 26) obtained using a multiple volume regression approach (Moran et al., 1999) of water collected with Niskin bottles from the CTD-rosette system within the ML (campaign-wide correction, method similar to Graff et al., 2023). We found that 29 of the 423 PON values were below the detection limit. Those values were removed. Negative values were within the relative standard deviation (RSD) of replicate measurements and therefore set to 0 μg N L^−1^. The RSD of replicate measurements associated with each fraction ranged between 9 % and 12 % for POC and between 24 % and 32 % for PON (Supporting Information **Table S1**). These values substituted for the uncertainty of the measurements that lacked replicates (2 POC values and 3 PON values).

Total particulate carbon (TC) was measured like POC but without prior acidification. Only one filter per fraction was collected due to sample volume constraints. Filtered volumes were the same as for POC. After subtraction of POC from TC, 39 out of 168 PIC values were negative (23 %). These values were substituted with a 0 μg PIC L^−1^.

Concentrations of biogenic and lithogenic silica (bSi and lSi) were determined by filtering subsamples onto 0.6-μm pore size polycarbonate membrane filters (47 mm PC, Isopore, Millipore), which were frozen at sea in cryovials at −20°C, transported to shore frozen, dried at 60°C, and then stored at room temperature until analysis. Filters were digested in Teflon tubes by adding 4 mL of 0.2 N NaOH (95°C, 40 minutes), cooled immediately, neutralized by adding 1 mL of 1 M HCl and centrifuged (10 minutes, 2,500 rpm) to separate lSi from bSi (Tréguer et al., 1992). The concentrations of bSi and lSi were assessed spectrophotometrically via the molybdosilicic acid method (Strickland and Parsons, 1968, Tréguer et al., 1992).

Concentrations of TEP were determined colorimetrically on triplicate samples of 400 mL each. Subsamples were filtered onto 0.4-μm pore size polycarbonate filters (25 mm, Whatmann, UK), stained with Alcian blue, and stored frozen at −20°C until analysis. Filter blanks, prepared by staining and rinsing wet filters, were processed like samples. Stained filters were soaked for at least 2 hours in 80 % sulfuric acid (95 % w/w; Fisher Scientific), and the absorption at 787 nm was measured spectrophotometrically (Passow and Alldredge, 1995). The stained filters were compared to a standard curve developed using xanthan gum (Sigma-Aldrich), and hence TEP determinations were expressed as standardized xanthan gum equivalents (XGeq; Bittar et al., 2018). The Alcian blue calibration f-factor was 34. For significant values of TEP to be determined the absorbance value of a sample at 787 nm must be at least twice the absorbance value of the blank at 787 nm (Passow and Alldredge, 1995). Ratios of TEP-C-to-POC were based on estimates of the carbon concentration of TEP using a conversion factor of 0.75 μg TEP-C per 1 μg XGeq TEP (Engel and Passow, 2001).

### Calculations of particle concentrations

The concentrations of suspended and sinking particles were calculated from the measured concentrations of *top*, *base,* and *tray* particle fractions by subtracting the *top* fraction from the *base* fraction and the *base* fraction from the *tray* fraction. The differences were scaled for the volume ratios as follows:

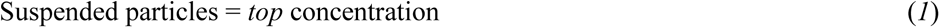

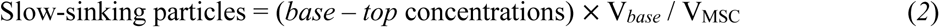

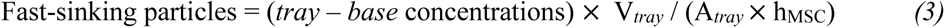

The volume of the base fraction (*V_base_*) associated with each deployment was calculated as:

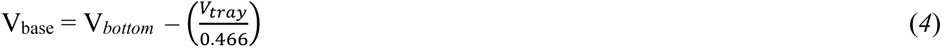

where V_MSC_ = 89.8 L, *V_bottom_* = 7.6 L, *V_tray_* was the volume of seawater we collected in the tray at each deployment (between 0.7 and 1.2 L) and 0.466 is a scaling factor obtained by accounting for the fact that the area of the tray covers only 46.6 % of the area of the bottom of the MSC. *V_base_* ranged between 5.1 and 6.1 L.

At 9 stations, fast-sinking marine snow (ESD >0.1 mm) particles were removed from the *tray* fraction before biochemical analysis. The remaining particles in the tray were smaller than 0.1 mm (ESD). This separation allowed us to assess the concentration of small fast-sinking particles (ESD <0.1 mm) separately from larger fast-sinking marine snow (ESD >0.1 mm). During sampling, we did not observe any marine snow in the *base* fraction, therefore we can assume that on average the slow-sinking particle population was smaller than 0.1 mm (ESD). Hence, for data interpretation, we summed the concentrations of slow-sinking particles and the concentrations of small, fast-sinking particles into a single particle fraction termed “small sinking particles” (**Table 2**).

**Table 2.**
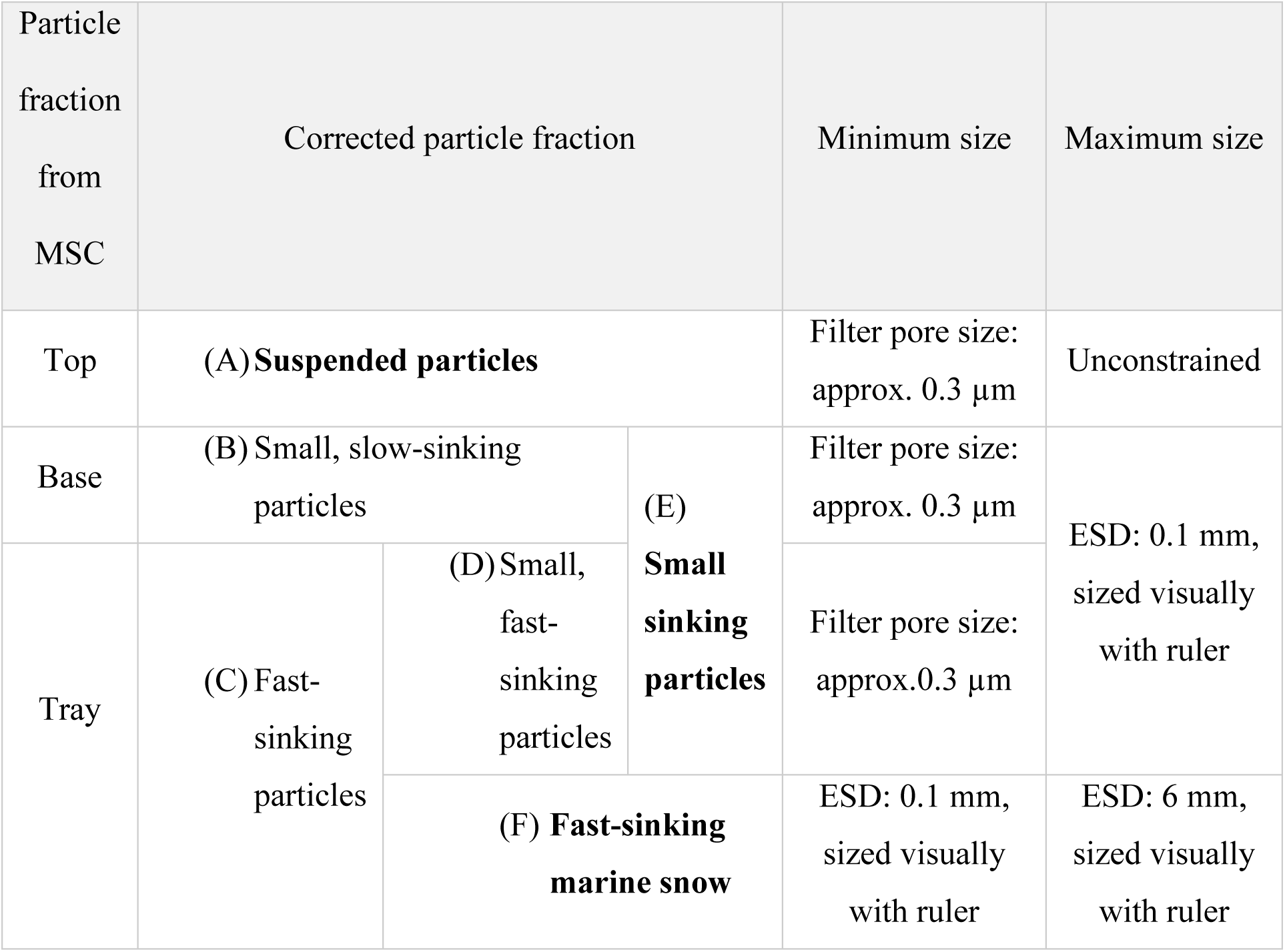
The nomenclature of distinct particle fractions sampled with the MSC and characterized in the present work. Bold text refers to the three particle fractions described in the results section. Letters (A)-(F) are used to identify particle fractions.

The concentration of fast-sinking marine snow present in the water column was calculated by dividing the number of marine snow particles present in each tray by the associated *V_tray_* and multiplied by the volume ratio 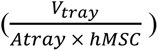. The volumetric concentration (mm^3^ L^−1^) of fast-sinking marine snow was calculated assuming the volume of a sphere and an average diameter of 0.5 mm, 1.5 mm, 3 mm, and 6 mm (midpoint of the 4 size bins: 0.1–1 mm, 1–2 mm, 2–4 mm, 4–8 mm).

In the Results section we describe the spatiotemporal variability of the following three particle fractions: suspended particles, small sinking particles, and fast-sinking marine snow (**Table 2**).

### POC concentrations associated with fast-sinking marine snow

We estimated the POC and PON mass (μg) associated with fast-sinking marine snow collected from MSC trays on May 27 at 60 m (below the MLD, 30 m). We placed 10–24 marine snow particles (approx. 1 mm, ± 10 %) on GF/F filters (n = 8) and measured the POC and PON mass as described below. The POC and PON mass (μg) of a single fast-sinking marine snow particle was calculated by dividing the blank-corrected POC and PON mass by the number of marine snow aggregates per filter. The dissolved organic carbon contained in the marine snow porewater (Alldredge, 2000) was likely lost during filtration. Thus, the POC mass is likely an underestimation of the total organic carbon transported by marine snow. The POC and PON mass (μg) associated with 2.5 mL of tray water was subtracted from the measured mass to account for contamination with smaller particles during pipetting. Five out of 8 values of PON were below the detection limit of the analytical instrument. The average POC mass measured on 1 mm marine snow particles was 0.7 ± 0.1 µg agg^−1^ (**Supporting Information Table S2**). This value is consistent with the one obtained using the fractal relationship of Alldredge (1998): POC (µg C) = 0.99 V^0.52^ (V is volume in mm^3^) for aggregate with an ESD of 1 mm. We hence used this fractal relationship to calculate the amount of POC in the collected fast-sinking marine snow particles with an average diameter of 0.5, 1.5, 3, and 6-mm and used a ± 25 % uncertainty, which account for the standard deviation of the replicated measurements of the POC mass of marine snow (15 %) and the error associated with the marine snow sizing (10 %).

Concentrations of POC associated with fast-sinking marine snow were obtained by multiplying the calculated POC mass (μg C) by the concentration of fast-sinking marine snow in each size bin.

To validate our method, we compared the POC concentrations directly measured in the particle fractions B “slow-sinking” and C “fast-sinking” (B+C) to the POC concentrations calculated by summing the concentrations of POC in E “small sinking” and F “fast-sinking sinking marine snow” so that B+C = E+F (see **Table 2** for nomenclature). The percent difference between the POC concentrations in “B+C” and “E+F” measured in adjacent stations was on average 15 % (n = 9), which is less than the standard deviation of the POC concentrations. This internal consistency in our calculations indicates that our method is appropriate.

A visual assessment of the particle morphology and phytoplankton community composition associated with fast-sinking marine snow was qualitatively performed on the 4 % formaldehyde-fixed samples collected on May 12, 17, 26 and 29 at 50–60 m depth using an Echo Revolve Microscope (Echo Laboratories, USA) with a 10x objective.

### Particle sinking velocity and POC fluxes

Particle fluxes were calculated by multiplying the concentration of sinking particles by an estimate of their average sinking velocity (Giering et al., 2016). Errors associated with the POC flux values were obtained by propagating the standard deviation of the measurements through each calculation. The sinking velocity of slow-sinking particles collected with the MSCs is on average 18 m d^−1^ based on the dimensions of the MSC (Giering et al., 2016), and the sinking velocity of fast-sinking particles is greater than 18 m d^−1^. This method does not allow further constraint of the sinking velocity of fast-sinking particles. We therefore used a novel approach that involves the co-deployment of MSCs and drifting sediment traps to estimate the sinking velocity of fast-sinking particles (small, fast-sinking particles + fast-sinking marine snow) assuming that sediment traps collect both slow-sinking and fast-sinking particles. There is evidence that traps may partially underestimate small particles due to hydrodynamics (Buesseler et al., 2007), therefore our sinking velocity and POC fluxes should be considered a lower bound. Two types of drifting sediment traps (a surface-tethered, drifting mooring and neutrally buoyant sediment traps) were deployed three times during the field campaign as described by Estapa et al. (2023). Methods described by Estapa et al. (2023) were used to measure bulk POC fluxes.

Three times (May 12, May 17, and May 25), we deployed the MSCs at the same depths as the drifting traps ((a) 73 m and 174 m, (b) 175 m, and (c) 95 m, 145 m, 195 m, and 500 m, respectively). To calculate the average sinking velocity of fast-sinking particles, we first calculated the POC flux due to fast-sinking particles measured by traps by subtracting the MSC POC flux of slow-sinking particles from the total POC fluxes measured with traps. Then we divided those fast-sinking POC fluxes by the POC concentrations measured in the MSC fast-sinking particle fraction, to give the sinking velocities of the fast-sinking particle fraction. This approach assumes that sinking velocities were near constant during the trap deployment period. These calculated sinking velocities of the fast-sinking particle fraction were used to calculate POC fluxes from the MSCs. The error associated with each sinking velocity and POC flux was calculated by propagating the standard deviation of the measurements through each calculation. As the sinking velocity of fast-sinking particles was assessed only on three days and at a few depths, we averaged the sinking velocities to be able to calculate the POC fluxes of particles collected in between such values (**Supporting Information Table S3**).

### Turbulent kinetic energy dissipation rates

Estimates of turbulent kinetic energy dissipation rates (*ϵ*, W kg^−1^) were derived from established scaling relationships (Lombardo and Gregg, 1989; D’Asaro, 2014) using air-sea momentum and buoyancy flux determinations calculated from bulk formulae (Johnson et al. 2023). These scaling estimates of *ϵ* are valid in the near-surface mixed layer.

### Statistical analyses

Simple linear regression was used to test (Student’s t-test, 95% confidence) for statistically significant temporal trends displayed by particle concentrations and biochemical concentrations. The linear analyses were performed using the function fitlm in Matlab (R2023a). The data presented in this study, and all the data generated during the EXPORTS field campaigns, can be found at NASA SeaBASS data repository.

## Results

### Turbulent kinetic energy dissipation rates in the mixed layer

The four storms created intense turbulence in the mixed layer where turbulent kinetic energy dissipation rates were surface intensified and varied in time by several orders of magnitude (**Figure 1**). The five sampling periods (P1 to P5) were characterized by turbulent kinetic energy dissipation rates of <10^−6^ W kg^−1^ in the mixed layer and were separated by intense storm events during which near-surface turbulent kinetic energy dissipation rates were >10^−4^ W kg^−1^ (**Figure 1**; D. A. Siegel et al., unpubl.).

### Concentrations of POC and TEP in the mixed layer

Particulate organic carbon (POC) concentrations measured at a nominal depth of ~20 m within the mixed layer depth decreased from 17±2 µmol C L^−1^ to 12±1 µmol C L^−1^ (n = 6) between the start and end of the field campaign (P2 and P5, respectively). Specifically, POC concentrations remained approximately constant between P2 and P3 and decreased by 1.4-fold from P3 to P5 (**Figure 2a**). The reduction in POC concentrations mirrored the decline in concentrations of Chlorophyll *a* and bSi in the mixed layer (Johnson et al., 2023; Meyers et al. 2023). The corresponding concentrations of TEP (**Figure 2b**) increased from 35±1 µg XGeq L^−1^ during P2 (n = 1, May 11) to 51±6 µg XGeq L^−1^ in P3 (n = 1, May 17), remained constant between P3 and P4, and decreased to 30±1 µg XGeq L^−1^ (n = 1) by the end of P5 (May 29). Concurrently, the relative contribution of TEP-C to POC (TEP-C-to-POC ratios) increased from 0.13±0.01 during P2 (n = 1, May 11) to 0.25±0.02 at the beginning of P5 (n = 1, May 24) (**Figure 2c**). The steepest decline in TEP and TEP-C-to-POC ratios occurred during P5.

**Figure 2.**
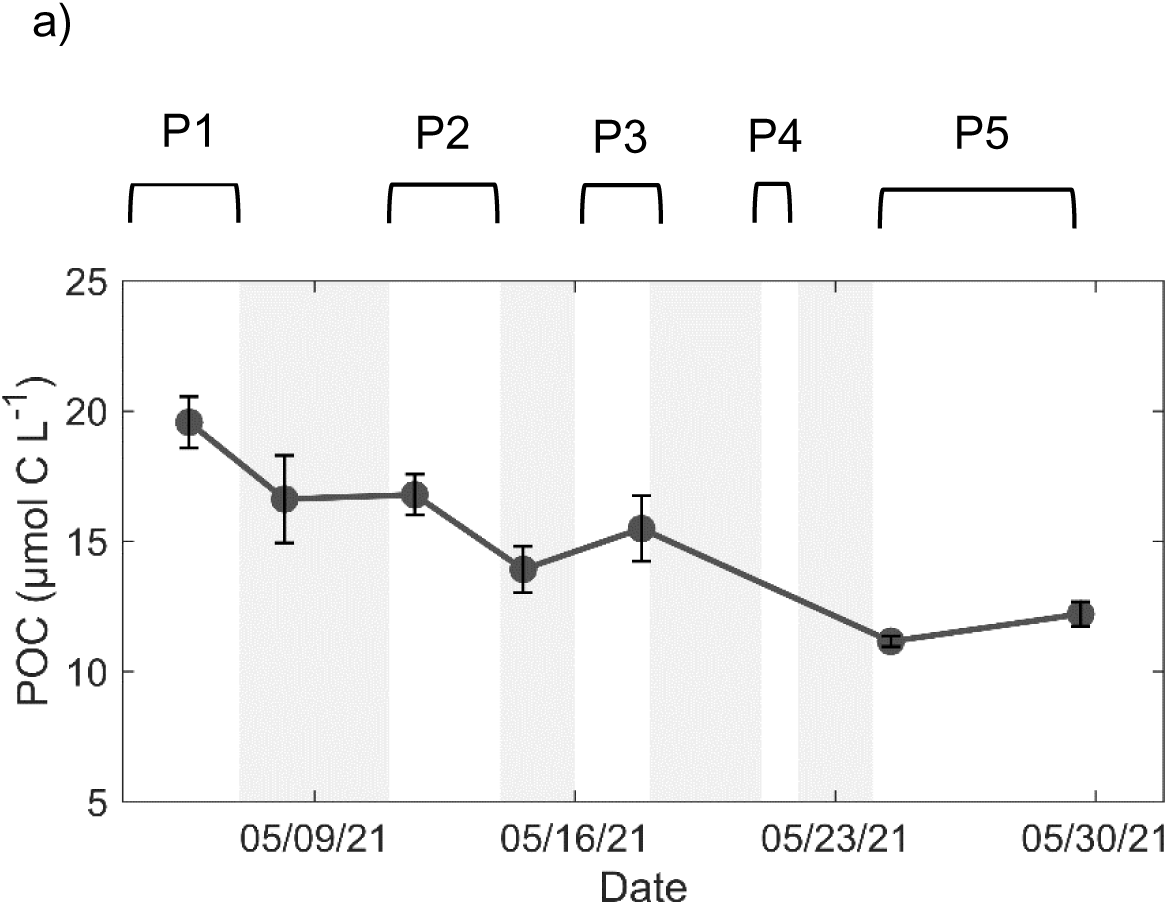
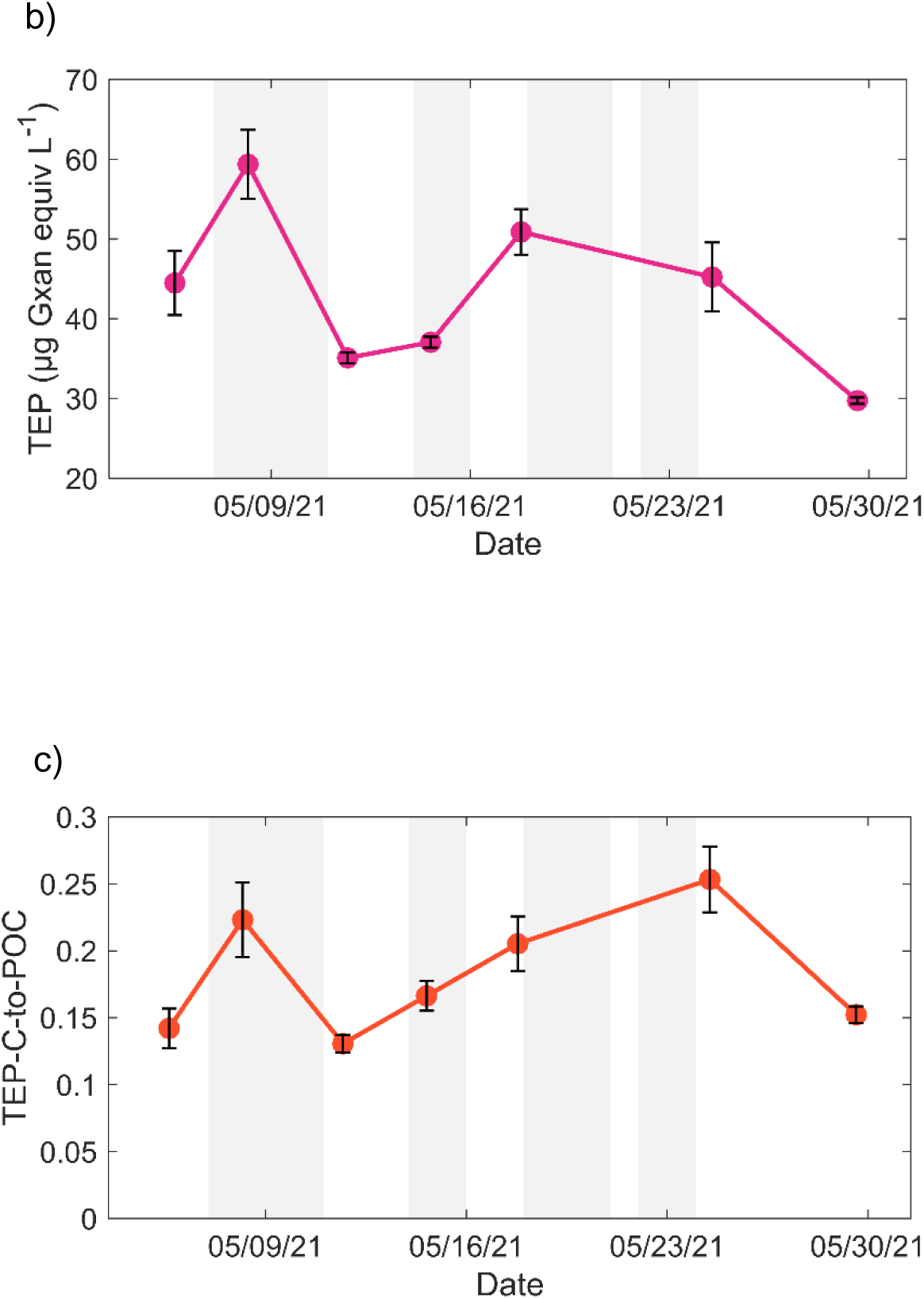
Change in the concentration of (a) POC and (b) TEP, and (c) TEP-C-to-POC ratios over time in seawater collected with Niskin bottles within the mixed layer at ~ 20 m depth. Note that the observations taken before May 8 are not included in our temporal analysis because 73 % of the mixed layer water was exchanged by the first storm. Gray shading represents the storms, which made sampling impossible. The five time periods demarcated by the storms (P1: May 4–7; P2: May 12–14; P3: May 16–19; P4: May 22; P5: May 25–29) are represented above panel a).

### Characteristics of suspended particles below the mixed layer

The POC concentrations of suspended particles collected with the marine snow catchers below the mixed layer at 50–75 m were highest after the first storm in P2 (May 13, 10±0.7 µmol C L^−1^, n = 1) and the second storm in P3 (May 17, 9.3±0.02 µmol C L^−1^, n = 1) (**Figure 3a**). Thereafter, POC of suspended particles at that depth decreased significantly from P3 to P5 (May 17–29; R^2^ = 0.61, p < 0.05, n = 8). Yet, they remained constant at 90–140 m, 330 m, and 500 m depth (no significant trend according to linear regressions; R^2^≤0.30, p ≥0.21, n = 10, 3, and 5, respectively) (**Figure 3a**). The PON and bSi concentrations of suspended particles broadly mirrored the spatiotemporal trend of suspended POC (Supporting Information **Fig. S2a, b**); however, the ratio between bSi and POC changed appreciably (**Figure 3b**). The molar bSi-to-POC ratio of suspended particles was the highest in P3 and decreased significantly from P3 to P5 at 50–75 m, 90–140 m and 330 m (R^2^ ≥0.70, p < 0.05, n = 6, 8 and 3, respectively). No significant temporal trend in the molar bSi-to-POC ratio of suspended particles was observed at 500 m depth (R^2^ = 0.36, p = 0.28, n = 5).

**Figure 3.**
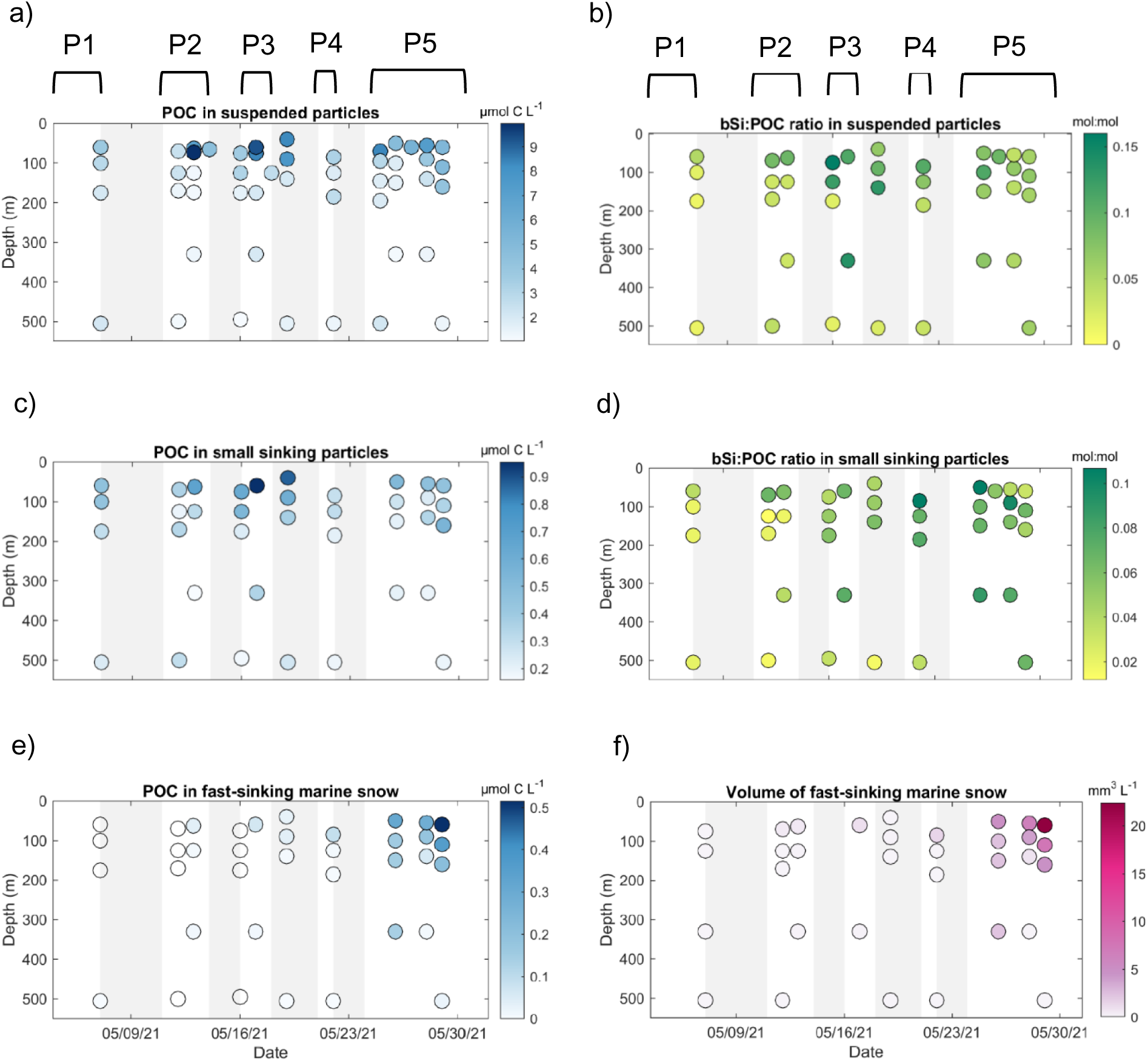
Time-depth distributions of particle characteristics for **(a-b)** suspended particles, **(c-d)** small sinking particles and **(e-f)** fast-sinking marine snow. The left-hand side panels show POC concentrations (a, c, e). The right-hand side panels show molar bSi-to-POC ratios for suspended and small sinking particles (b, d) and, volumetric concentrations of fast-sinking marine snow particles (f). The five time periods (P) demarcated by the storms (P1: May 4–7; P2: May 12–14; P3: May 16–19; P4: May 22; P5: May 25–29) are represented above panels a) and b).

### Characteristics of small sinking particles below the mixed layer

The POC concentration of small sinking particles peaked at 50–75 m during P3 (May 17) (0.95±0.05 µmol C L^−1^, n = 1) and decreased significantly from P3 to P5 (R^2^ = 0.94, p <0.007, n = 5; **Figure 3c**). The POC concentrations of small sinking particles did not show a significant change over time below the 50–75 m depth layer (R^2^ ≤0.29, p ≥0.21, n = 10, 4 and 7 at 90–140 m, 330 m, and 500 m, respectively). The concentration of PON in small sinking particles mirrored the evolution of POC (Supporting Information **Fig. S2c**).

The bSi concentration of small sinking particles at 50–75 m was the highest during P3 (May 17) and at the beginning of P5 (May 26). Between 90 m and 330 m depth, bSi concentrations increased from P2 to P3, whereas at 500 m, a significant increase was observed from P4 to P5 (R^2^ = 0.74, p = 0.05, n = 5) (Supporting Information **Fig. S2d**).

The molar ratios of bSi-to-POC of small sinking particles at 50–75 m depth doubled from P3 to the beginning of P5 (May 26) and halved within P5 from May 26 to May 29 (**Figure 3d**). At 90–140 m and 500 m depth, the bSi-POC ratios increased significantly from P2 to P5 (respectively: R^2^ = 0.70, p < 0.003, n = 10, and R^2^ = 0.5, p = 0.05, n = 5).

Molar PIC-to-POC ratios in small sinking particles did not show any significant temporal evolution throughout the field campaign (R^2^≤0.43, p = ≤0.77, n = 8, 8, 4 and 5 at 50–75 m, 90– 140 m, 330 m, and 500 m, respectively) (Supporting Information **Fig. S2e**). Similarly, molar POC-to-PON ratios did not show any spatiotemporal trend over time or depth. However, molar lSi-to-POC ratios of small sinking particles (n = 3) showed a sharp decrease over time due to the complete disappearance of lSi in small sinking particles at the end of P5 (May 28) (Supporting Information **Fig. S2f**).

Taken together (**Fig. 3b, d**) the changes of bSi:POC in suspended and small sinking particles over time and depths suggest sinking of small, silica-rich particles, which in the discussion, we refer to as the first sedimentation episode.

### Characteristics of fast-sinking marine snow below the mixed layer

The volumetric concentration (mm^3^ L^−1^) of fast-sinking marine snow increased exponentially from P2 to P5 (R^2^= 0.95–0.99, n = 7, 8 and 5 at 50–75 m, 90–125 m and 500 m, respectively) with P5 characterized by the highest volume concentrations of the field campaign (**Figure 3f**). In agreement with the increase in marine snow volumes, the POC concentrations of fast-sinking marine snow increased significantly over time from P2 to P5 (R^2^ = 0.66–0.81, p < 0.05 n = 6 and 6 at 50–75 m and 90–125 m, respectively) and were highest at the end of P5 (May 29), when they were 0.52 µmol C L^−1^ and 0.02 µmol C L^−1^ at 60 m and 500 m, respectively (**Figure 3e**). In the discussion we refer to the sinking of fast-sinking marine snow aggregates as the second sedimentation episode.

The plankton community composition associated with fast-sinking marine snow collected at 50–75 m transitioned during the field campaign. Overall, we observed the presence of detrital diatoms initially and an increased contribution of heterotrophs and healthy diatoms by the end of P5. Specifically, in P2 (May 12), fast-sinking marine snow contained mostly detrital diatoms (empty and broken frustules) mainly *Chaetoceros* spp. and Navicula-type pennates, but also *Rhizosolenia* spp., silicoflagellates (haptophytes), and Tintinnids (heterotrophic ciliates). In P3 (May 17), we observed an increase in the relative abundance of dinoflagellates, including autotrophs (Ceratium spp.) and hetero- or mixotrophic taxa, whereas the relative abundance of silicoflagellates and *Chaetoceros* spp. appeared to be decreased compared to May 12. At the beginning of P5 (May 26), marine snow composition was similar to the one observed in P3 with the addition of radiolarians. On the last day of P5 (May 29), marine snow did not contain silicoflagellates and was likely largely composed of heterotrophs (dinoflagellates, tintinnids): although intact diatoms also increased (Supporting Information **Fig. S3**).

### Sinking velocity of fast-sinking particles

The sinking velocity of fast-sinking particles (small fast-sinking particles plus fast-sinking marine snow) above 200 m depth increased over time from an average (± variability among estimates) of 17±5 m d^−1^ (n = 5) during P2 and P3 to an average of 97±30 m d^−1^ (n = 4) during P5 (**Figure 4**, **Supporting Information Table S3**). This increase was concurrent with the increase in the volumetric concentration of fast-sinking marine snow observed during P5 (**Figure 2f**). From P2 to P5, the contribution of small silica-rich particles to the total POC content of sinking particles decreased from, on average, 99 % to 54 %. This indicates that sinking velocities estimated during P2 and P3 were almost fully representative of the sinking velocity of small silica-rich particles, whereas the sinking velocity during P5 includes the contribution of fast-sinking marine snow. During P2 and P3, when marine snow was rare, fast-sinking particles sank at approximately the same velocity as slow-sinking particles (18 m d^−1^). At the beginning of P5 at 500 m depth, the sinking velocity of fast-sinking particles was 42 ± 5 m d^−1^ (n = 1) (**Figure 4**).

**Figure 4.**
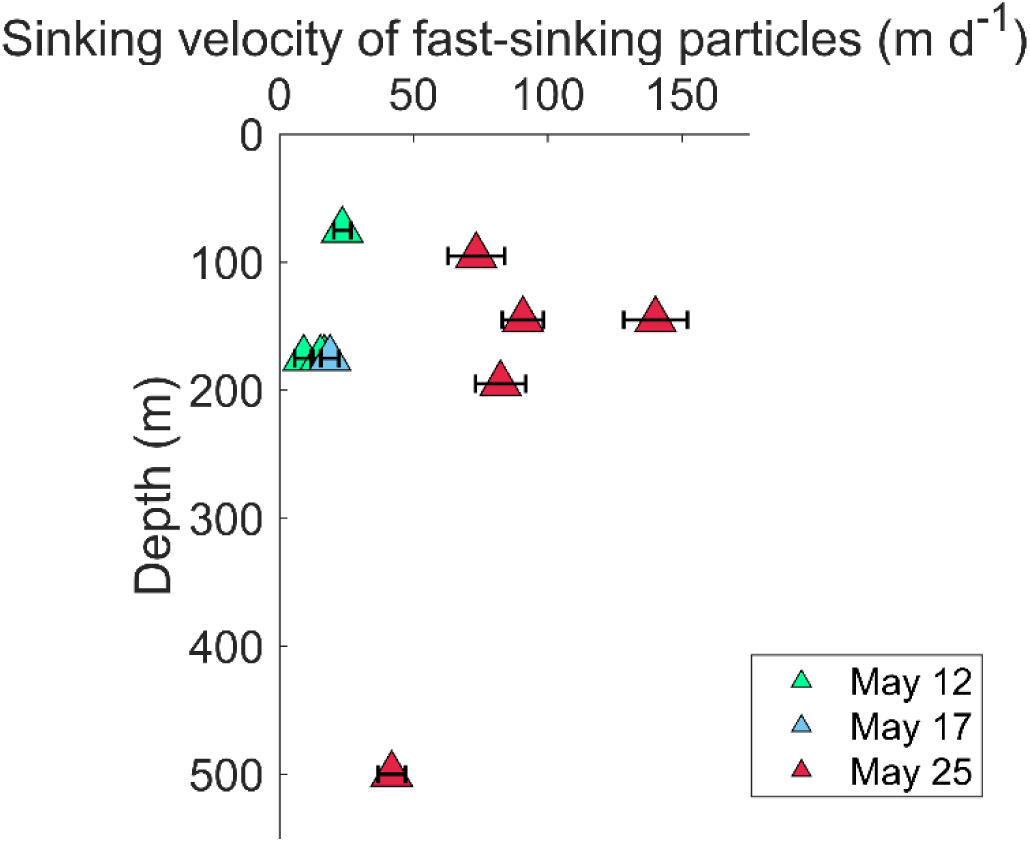
Temporal and vertical variations in the estimated sinking velocities of fast-sinking particles (small, fast-sinking particles plus fast-sinking marine snow) calculated from a comparison of MSC collections and bulk sediment trap POC fluxes. Error bars are the propagated measurement standard deviations.

### POC fluxes

The POC flux of the total sinking particle pool (slow-sinking plus fast-sinking particles) peaked after the fourth storm at the end of P5 (May 29). Overall, values increased significantly over time from P2 to P5 at the depths 50–75 m and 90–140 m (R^2^ = 0.58–0.87, p < 0.001, n = 13 and 14, respectively) and, although not significantly, at 500 m (R^2^ = 0.36, p = 0.15, n = 7) (**Figure 5a**). Specifically, at 90–125 m fluxes were 6±3 mmol C m^−2^ d^−1^ (n = 8) during P2–4 and a factor of 4.7 higher (30±12 mmol C m^−2^ d^−1^) during P5 (n = 6). At 500 m depth, fluxes increased from P2–4 to P5: from 4±1 mmol C m^−2^ d^−1^ (n = 4) to 6±1 mmol C m^−2^ d^−1^ (n = 2).

**Figure 5.**
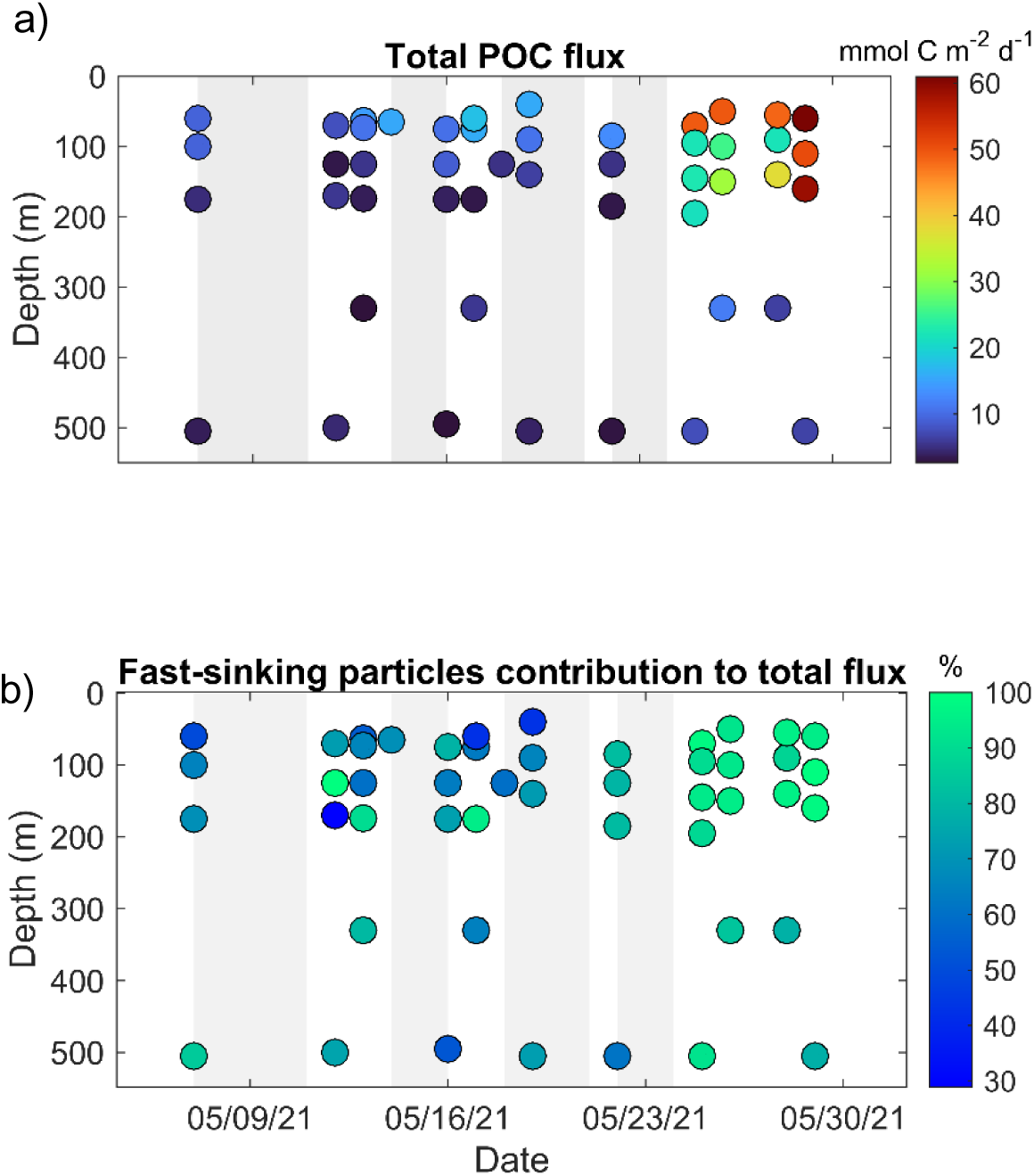
Time-depth distributions of (a) total POC flux (due to both slow-sinking and fast-sinking particles) and (b) fractional (%) contribution of POC flux due to fast-sinking particles to total POC flux.

The contribution of fast-sinking particles to the total POC flux (slow-sinking plus fast-sinking particles) was highest (95 %) during P5 between 50–70 m depth. At 50–75 m and 90– 125 m depth, the contribution of fast-sinking particles to the total POC flux increased significantly from P2 to P5 (between May 13 and May 29; R^2^ ≥0.67, p < 0.003, n = 9 and 10, respectively; **Figure 5b**). No significant temporal trend was observed at 500 m (R^2^ = 0.004, p = 0.90, n = 6).

## Discussion

The EXPORTS North Atlantic field sampling started at the end of a bloom of diatoms when 72 % of the bSi produced during the bloom had already been exported (M. A. Brzezinski et al., unpubl.). The extremely low silicate (SiO_4_ ~ 0.2 µmol L^−1^) and high nitrate (NO_3_ ~ 5 µmol L^−1^) concentrations that we observed in the mixed layer at the start of the field campaign suggest that silicate limitation was the likely trigger of the diatom bloom decline (Johnson et al., 2023; M. A. Brzezinski et al., unpubl.). Consistent with this interpretation, the phytoplankton community in the mixed layer was dominated by diatoms at the beginning of the study (P1) but their absolute and relative abundances declined throughout the field campaign (Meyer et al., 2023). An appreciable shift in the phytoplankton community from diatoms to haptophytes and chlorophytes in the mixed layer during the field campaign (Meyer et al., 2023; E. Jones, pers. comm.) indicated that the phytoplankton bloom was likely transitioning because of silica limitation (Sieracki et al., 1993). Nevertheless, diatom contribution to POC export during the field campaign was estimated to be 40–70 %, partially due the rapid use by diatoms of SiO_4_ entrained into the mixed layer by the storms (M. A. Brzezinski et al., unpubl.).

We observed two sedimentation episodes that followed the main diatom bloom sedimentation that occurred prior to the ship-based observations (**Figure 6**). The first sedimentation episode (dominating flux from P1 to P4) was dominated by small, silica-rich particles that sank in the upper mesopelagic during the storm periods. The second sedimentation episode (predominantly observed during P5) consisted of fast-sinking marine snow aggregates, whose volume increased after the storms ceased. Our data indicate that these two sedimentation episodes (sinking diatoms and sinking marine snow) likely overlapped. During P4, 24 % of the POC concentration measured in sinking particles was due to fast-sinking marine snow at 85 m depth; however, between 125 m and 500 m depth, sinking diatoms were still dominating (99 % of POC in sinking particles). By the beginning of P5 (May 25) the contribution of fast-sinking marine snow increased to 39 % at 50–330 m. This comparison indicates that the contribution of marine snow to the POC flux was increasing from P4 to P5. The overlap of the two sedimentation episodes is also supported by drifting sediment traps data that show increased fluxes of biogenic silica, aggregates and POC in P4 (traps were deployed on May 22–24, Estapa et al., 2023). Different depths of the water column were characterized by different particle dynamics indicating the concurrent presence of multiple processes that affect the transfer of POC through the water column.

**Figure 6.**
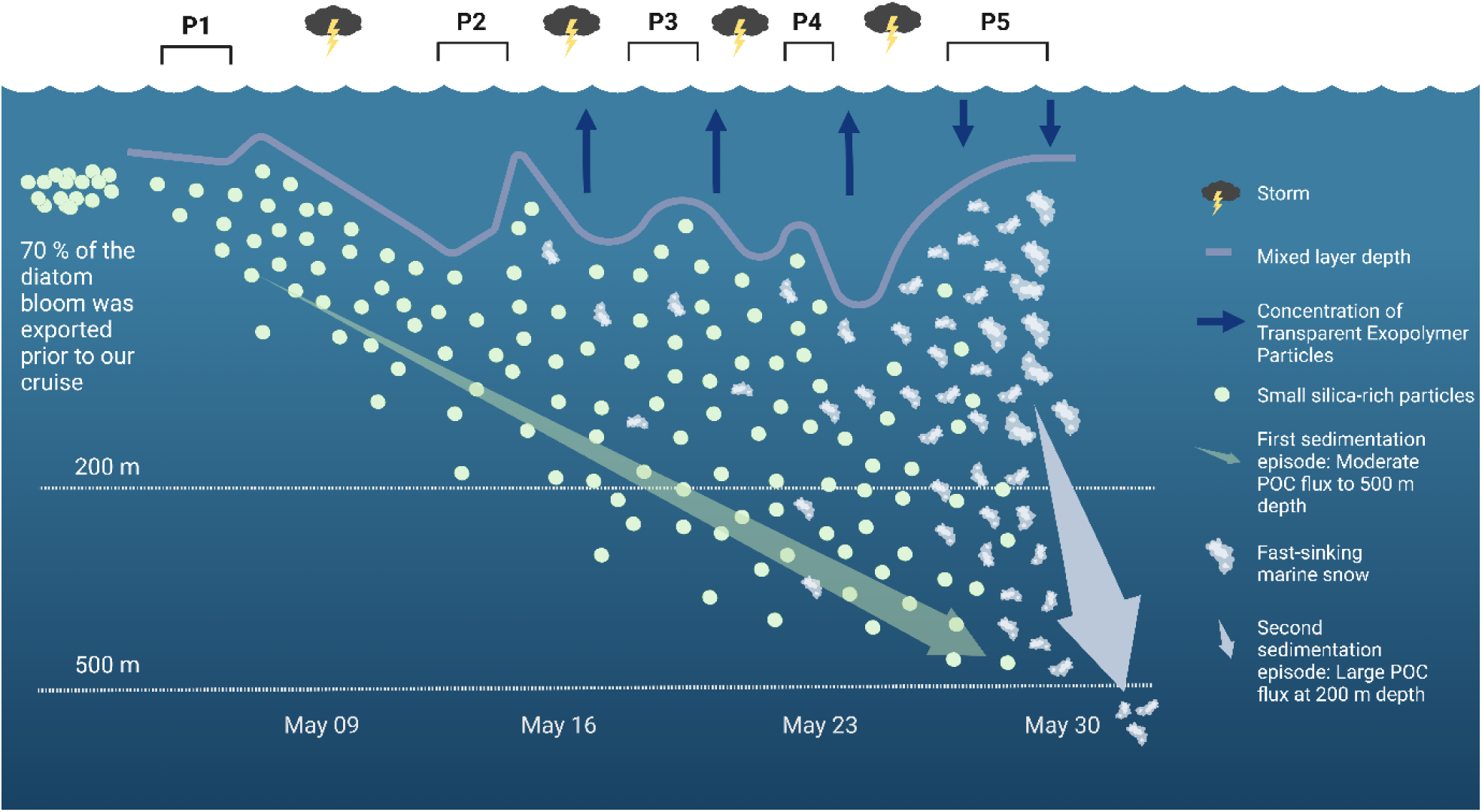
Schematic of the two sedimentation episodes, which followed the main diatom export prior to our arrival on this site. **Pre-field campaign** Nutrient and biomass analysis revealed that a large diatom bloom had occurred and mostly sank before our visit (Johnson et al., 2023; M. A. Brzezinski et al., unpubl.). **P2-P4** First sedimentation episode (transparent green arrow): Small, silica-rich particles sank in the upper mesopelagic during the period characterized by the four intense storms. Concurrently, TEP accumulated in the mixed layer and the volume of fast-sinking marine snow remained low below the mixed layer. Small, silica-rich particles created a moderate POC flux at 500 m depth. **P5** Second sedimentation episode (transparent white arrow): The volume of fast-sinking marine snow increased exponentially below the mixed layer, while TEP concentrations decreased within the mixed layer. Marine snow appeared abundantly at the beginning of P5 when the storms ceased, turbulence was moderately intense, and the mixed layer depth decreased to approximately 30 m depth. This event was characterized by a large POC flux at 200 m depth. **Post-field campaign** The second sedimentation episode displayed the potential for substantial and efficient POC export at and below 500 m depth. The schematic was created using ©BioRender.

In the following sections, we characterize each sedimentation episode and discuss each one’s contribution to POC transport in the upper mesopelagic.

### Moderate POC fluxes due to small sinking particles during the storms

Our data, particularly the molar bSi-to-POC ratios, indicate that the flux of small silica-rich particles below the mixed layer increased during the storm periods (P2–P4) and reached 500 m depth. These observations are consistent with the increased flux of intact and empty and fragmented sinking diatom cells collected in polyacrylamide gel sediment traps between P2 and P4 (Bodel, 2023). We further observed an accumulation of TEP in the mixed layer during this period, indicating a faster production of TEP than loss of TEP from, for example, export or consumption. Enhanced TEP production has been linked to stressed and senescent phytoplankton, which lends further support to the idea that we sampled the termination phase of a diatom bloom (Passow and Alldredge, 1995). We expected the combination of senescent diatoms, increased TEP concentrations and elevated particle collision rates due to high levels of mixed layer turbulence (**Figure 1**, see section to follow) to result in sinking of marine snow aggregates. Instead, while TEP was accumulating in the mixed layer during P2 to P4 (i.e., increased TEP-C-to-POC ratio), sinking fluxes were mostly dominated by small silica-rich particles (i.e., diatoms and their detritus).

We suggest that the presence of the four intense storms in succession led to the sinking of these small silica-rich particles. First, the high turbulent kinetic energy dissipation rates created by the four storms in the mixed layer (mean turbulent kinetic energy dissipation rates > 10^−6^ W kg^−1^, **Figure 1**) may have favored disaggregation of marine snow over aggregation (Takeuchi et al., 2019; D. A. Siegel et al., unpubl.). These storms may have limited the size of marine snow allowing smaller particles to sink below the mixed layer. In addition, the storm-driven deepening of the mixed layer redistributed particles in an additional 25–40 m of the water column (Johnson et al., 2023). Based on coagulation theory, lower particle concentrations (for example due to particle dilution in the water column) decrease particle collision rates limiting aggregation (Jackson et al., 1990). Hence, storms may have allowed TEP to form from TEP precursors exuded by senescent diatoms, but the concentration of particles was too low to allow the formation of aggregates. Finally, particle redistribution may have isolated surface particles below the mixed layer each time the mixed layer shallowed again, allowing them to sink deeper. A silica budget analysis performed by M. A. Brzezinski et al., (unpubl.) indicates that bSi isolated below the mixed layer in between storms may have accounted for a significant fraction of the bSi exported during P1–P3, but not during P4–P5, when sinking of marine snow aggregates likely controlled bSi export.

Our data on fluxes dominated by small silica-rich particles indicate that sinking diatoms were the main contributors of the POC flux during the storm periods. The sinking velocity of particles in water is a function of their size and density (e.g., Passow and De La Rocha, 2006; Cael et al., 2021). As the density of silica (2000 kg m^−3^) is higher than that of organic carbon, preferential sedimentation of silica-rich particles is consistent with expectations. These silica-rich particles were small (< 0.1 mm) and sinking slowly (~18 m d^−1^, **Figure 4**); hence, their POC content was likely mostly recycled through the food web in the shallow mesopelagic rather than transported to below 500 m. This shallow remineralization is supported by the lack of an increase in the POC concentrations of small sinking particles below 200 m depth (**Figure 3c**). Flux profiles estimated from Thorium-234 disequilibrium substantiate this idea, as they show that during P2–P4, POC fluxes decreased while bSi fluxes increased between the mixed layer and 100 m depth (Clevenger et al., 2023). Though we observed that during the cruise small sinking particles contributed on average 98 ± 5 % (n = 4) of the POC concentration of sinking particle population at 500 m depth, we cannot exclude the possibility that some of these small particles originated from disaggregation of diatom aggregates rather than sinking there directly. Disaggregation of larger particles is supported by an increased aggregate concentration observed in the upper mesopelagic (above 500 m depth) beginning on P4 as measured by the MSC and drifting sediment traps.

Overall, our observations suggest that small, silica-rich, slow-sinking particles, namely diatoms and their fragments can dominate the flux of POC below the mixed layer during the decline of a spring diatom bloom.

### High POC fluxes due to fast-sinking marine snow after the storms

The large and rapid sedimentation episode of fast-sinking marine snow aggregates mainly developed when the last storm ceased, more than three weeks after the diatom bloom. This observation challenges the standard view on diatom-aggregate export happening as a single event at the declining phase of the bloom. Furthermore, it suggests that intense storms inhibited the formation and or sinking of marine snow aggregates.

Between P2 and P5, intense winds and surface waves generated by four strong storms (≥ 9 on the Beaufort wind force scale) led to turbulent kinetic energy dissipation rates in the mixed layer larger than 10^−5^ W kg^−1^ (**Figure 1**) and to a deepening of the mixed layer depth by 25 m to 45 m (Johnson et al., 2023). Turbulence has been found to enhance marine snow formation up to dissipation rates of 10^−6^ W kg^−1^ (Takeuchi et al., 2019). However, for higher dissipation rates, disaggregation may outcompete aggregation and hence limit marine snow size (Alldredge et al., 1990; Takeuchi et al., 2019). After the last storm (P5), the rates of turbulent dissipation energy in the mixed layer decreased to values less than 10^−7^ W kg^−1^ (**Figure 1**) which favors particle aggregation (Takeuchi et al., 2019), and allowed, at least partially, the observed rapid increase of fast-sinking marine snow within and below the mixed layer (D. A. Siegel et al., unpubl.). Furthermore, the shoaling of the mixed layer to approximately 30 m during P5 likely allowed particles to concentrate, potentially increasing particle-particle collision rates. The formation of marine snow due to aggregation requires the presence of TEP which provides stickiness, promoting successful aggregation after collision (Logan et al., 1995). The observed decreased concentrations of mixed layer TEP in P5 is consistent with TEP sinking as part of the marine snow aggregates. It is therefore likely that the combination of moderate turbulence levels and a shallow ML depth triggered the rapid, exponential aggregation of fast-sinking marine snow that we observed in the eddy core waters during P5. Data obtained using the Underwater Visual Profiler (Picheral et al., 2015) during the field campaign indicates the formation of large particles and loss of small particles above 150 m depth during P4–P5 (D. A. Siegel et al., unpubl.) supporting our observations. Similarly, Giering et al., (2016) found a two orders-of-magnitude increase in the concentration of large particles above 100 m depth after a strong spring storm (9– 10 on the Beaufort wind force scale) in the high latitudes of the North Atlantic.

The consequence of a “postponed” formation of aggregates should result in the sinking of aggregates containing a more heterogeneous plankton community. Consistent with this expectation, the composition of fast-sinking marine snow aggregates changed over time and shifted from being dominated by detrital diatoms to a community that presented increased contributions of intact diatoms and heterotrophic taxa (i.e., dinoflagellates and tintinnids).The observed intact diatoms in sinking marine snow may have originated from growing diatoms that were using the silicic acid injected by the mixed layer entrainment caused by the storms (M. A. Brzezinski et al., unpubl.), as diatoms may recover from silica limitation within hours (De La Rocha and Passow, 2004). High concentrations of dinoflagellates and tintinnids have previously been observed at the end of diatom blooms, indicating that these heterotrophs may be significant consumers of senescent bloom-forming diatoms (Passow and Peinert, 1993; Tiselius and Kuylenstierna, 1996; Sherr and Sherr, 2007).

Our data indicates that in P5 fast-sinking marine snow created a large POC flux below the mixed layer. The calculated POC fluxes at 100 m depth (51±15 mmol C m^−2^ d^−1^) are indeed on the high end of observed upper ocean POC fluxes (Mouw et al., 2016; Bisson et al., 2020; Giering et al., 2023 spring bloom in South Georgia, 54 mmol C m^−2^ d^−1^, 95 m). In addition, our data suggests that such POC flux had the potential to rapidly reach the depth of the winter mixed layer (600 m in the North Atlantic, de Boyer Montégut et al., 2004) and hence to be sequestered for a relevant amount of time. This statement is supported by the large carbon content (0.7 µg C agg^−1^) of the marine snow aggregates and the high average sinking velocities of fast-sinking particles (97±30 m d^−1^, n = 3). The measured average sinking velocity represents an average of the entire fast-sinking particle population, including particles smaller than 0.1 mm in diameter. Hence the large marine snow that we observed was likely sinking faster than our estimate. In addition, the sinking velocity measured at the beginning of P5 (May 25) is likely an underestimation of the sinking velocity of the fast-sinking particles presents in the water at the end of P5 (May 29): In fact, the volumetric concentration of fast-sinking marine snow at 60 m depth increased by 3.5-fold during P5 increasing the proportion of marine snow to the total fast-sinking particle pool.

Overall, our observations indicate that storm-induced turbulence and mixed layer deepening created the conditions for the development of a marine snow sedimentation episode that allowed the sinking of POC, TEP-C and a transitioning, mixed plankton community, which likely would have not sunk otherwise.

## Conclusions

Our work demonstrates that observations from the MSCs complement the traditional suite of sampling devices (e.g. Niskin bottle, sediment traps and more recently UVP) allowing detailed observations of particle size and sinking velocity at relatively high temporal resolution. We found that a significant amount of the POC flux originated from a spring diatom bloom weeks after the main bloom sedimentation event. Moreover, the MSCs enabled us to observe the presence of two distinct sedimentation episodes that contributed differently to POC flux below the mixed layer and did not follow the commonly depicted export progression at the end of a spring bloom. Rather than the export of large, fast-sinking, diatom-dominated phytodetrital aggregates, we first observed the sinking of small particles (likely individual diatoms or small diatom-aggregates). The sinking of marine snow aggregates happened later, when the storms ceased, and contributed to the export of a more mixed plankton community. Our data suggests that the succession of intense storms may have been the key for this unorthodox flux progression. Changes in turbulent kinetic energy dissipation rates and in mixed layer depths likely determined not only the magnitude but also the composition of the flux of POC during this critical phase of annual export and carbon sequestration. Ultimately, our findings suggest that increased storm severity due to climate change (e.g., Knutson et al., 2010) may have important implications for the timing of POC export and the amount and type of carbon exported in the upper mesopelagic zone.

## Acknowledgements

The authors are grateful to the EXPORTS science team for the collaborative effort, data and ideas exchange and scientific production that acted as a baseline for the present work. A big thanks goes to the captain and crew of the RRS James Cook. We also thank Dr. Chelsey Baker and Jack Williams who analyzed the TEP samples. Finally, we thank Dr. Mark Brzezinski for the valuable discussions during the preparation of the manuscript.

## Data Availability Statement

All data presented here are publicly available on the NASA data repository, SeaBASS along with all other EXPORTS data. The data can be accessed here: https://seabass.gsfc.nasa.gov/experiment/EXPORTS.

## Supporting Information

### Supporting Information Tables

**Table S1:** The average relative standard deviation (RSD) of replicate measurements of POC and PON concentrations in each particle fraction collected with the Marine Snow Catcher.

| RSD (%) | T0 | top | base | tray |
| --- | --- | --- | --- | --- |
| POC | 9 | 8 | 12 | 10 |
| PON | 28 | 30 | 24 | 32 |

**Supporting Information Table S2:** The measured POC mass of fast-sinking marine snow of 1mm equivalent spherical diameter (ESD). Marine snow was collected at 60 m on May 27. Values marked as DL were below the detection limit (DL) of the analyzer.

| Filtered fast-sinking marine snow | Average POC | Average PON |
| --- | --- | --- |
| number | $\mu\text{g C aggregate}^{-1}$ | $\mu\text{g N aggregate}^{-1}$ |
| 20 | 0.64 | DL |
| 20 | 0.58 | 0.08 |
| 20 | 0.66 | 0.14 |
| 10 | 0.58 | DL |
| 20 | 0.67 | DL |
| 20 | 0.44 | DL |
| 24 | 0.74 | DL |
| 20 | 0.93 | 0.23 |
| **Average** | **0.65** | **0.07** |
| **SD** | **0.10** | **0.04** |

**Supporting Information Table S3.** Calculated average sinking velocities used to calculate POC fluxes due to fast-sinking particles.

| Depth (m) | P1–2 | P3 | P4 | P5 |
| --- | --- | --- | --- | --- |
| 40–100 | $23 \pm 3 \text{ m d}^{-1}$ | $17 \pm 5 \text{ m d}^{-1}$ | $43 \pm 43 \text{ m d}^{-1}$ | $73 \pm 10 \text{ m d}^{-1}$ |
| 125–150 | $17 \pm 5 \text{ m d}^{-1}$ | $17 \pm 5 \text{ m d}^{-1}$ | $17 \pm 5 \text{ m d}^{-1}$ | $115 \pm 35 \text{ m d}^{-1}$ |
| 175–500 | $15 \pm 4 \text{ m d}^{-1}$ | $15 \pm 4 \text{ m d}^{-1}$ | $15 \pm 4 \text{ m d}^{-1}$ | - |
| 160–195 | - | - | - | $82 \pm 9 \text{ m d}^{-1}$ |
| 300–500 | - | - | - | $42 \pm 5 \text{ m d}^{-1}$ |

### Supporting Information Figures

**Fig. S1.**
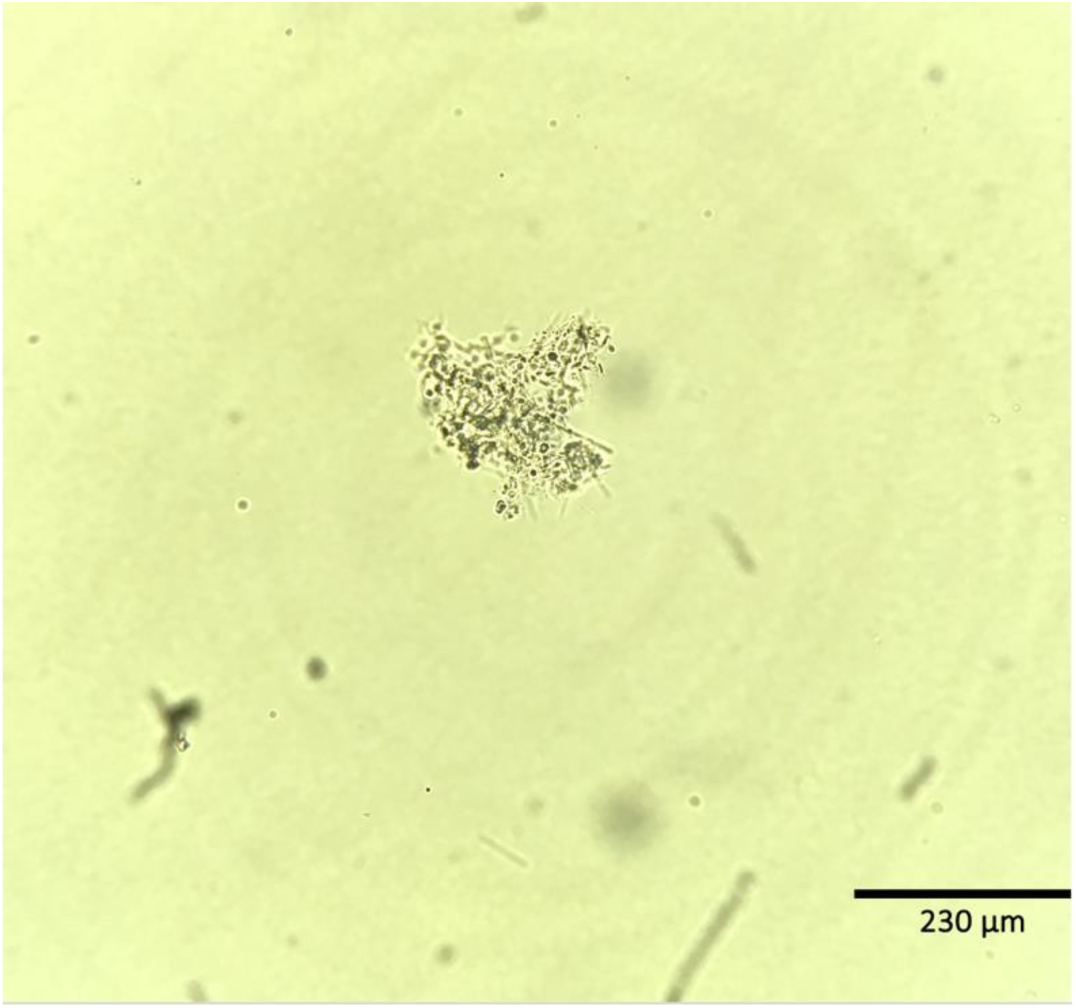
Example of a microscopy photo of intact fast-sinking marine snow (a) collected at 50 m on May 29.

**Fig. S2.**
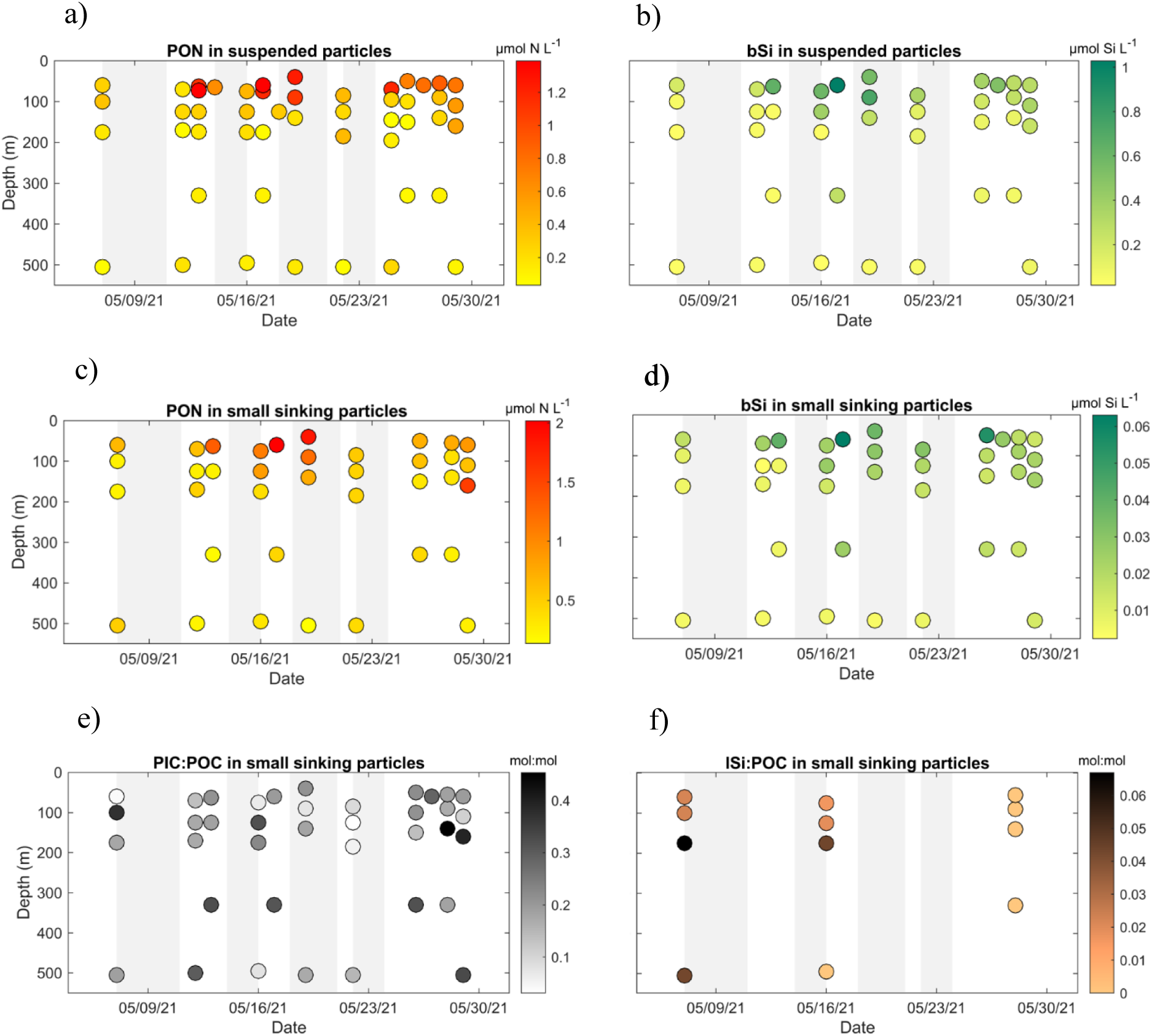
Top panels: Temporal change of (a) POC (b) bSi concentrations in suspended particles between 40 and 500 m depth. Middle panels: Temporal change of (c) POC (d) bSi concentrations in small sinking particles between 40 and 500 m depth. Bottom panels: Temporal change of (e) molar PIC-to-POC ratios and (f) molar lSi-to-POC ratios in small sinking particles between 40 and 500 m depth.

**Fig. S3.**
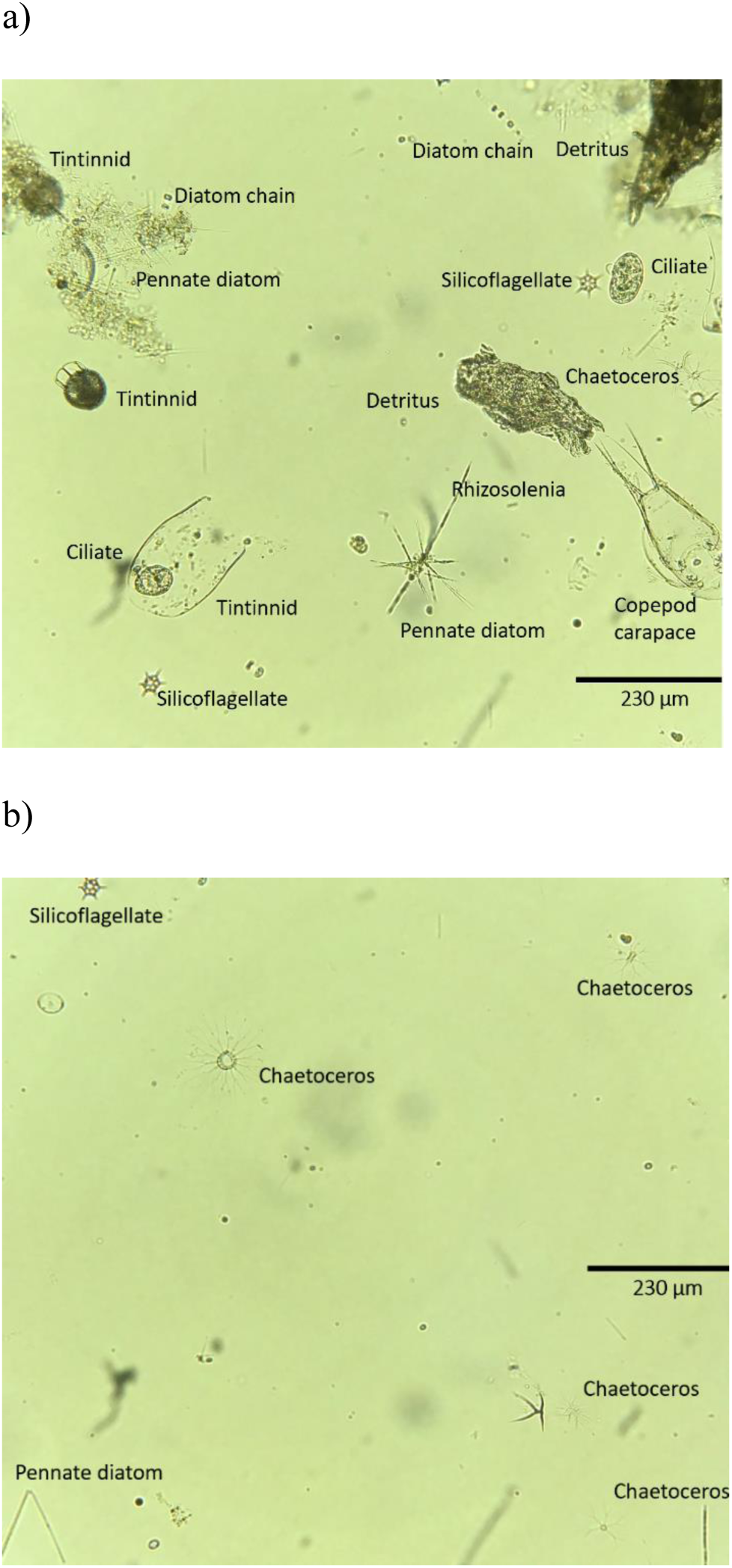
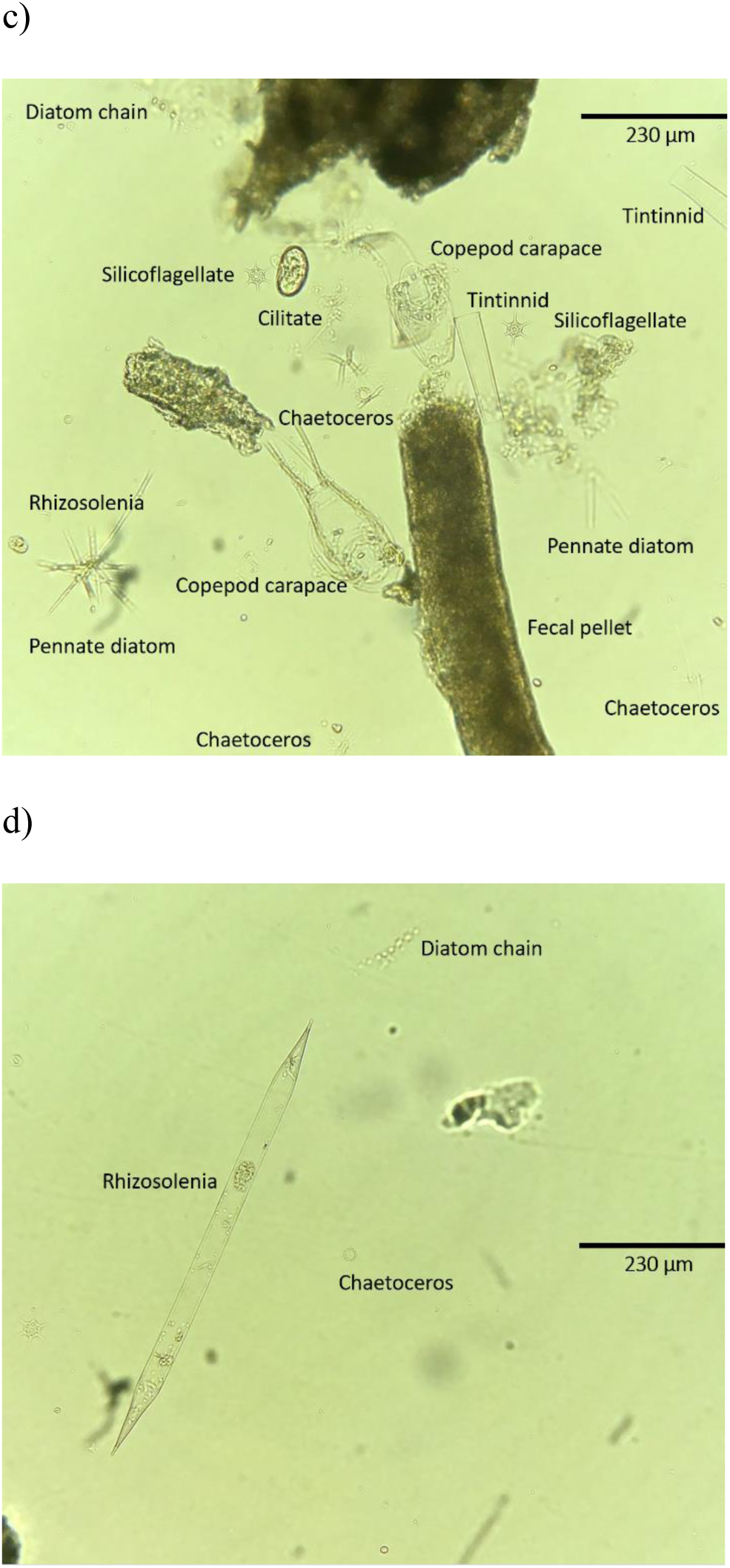
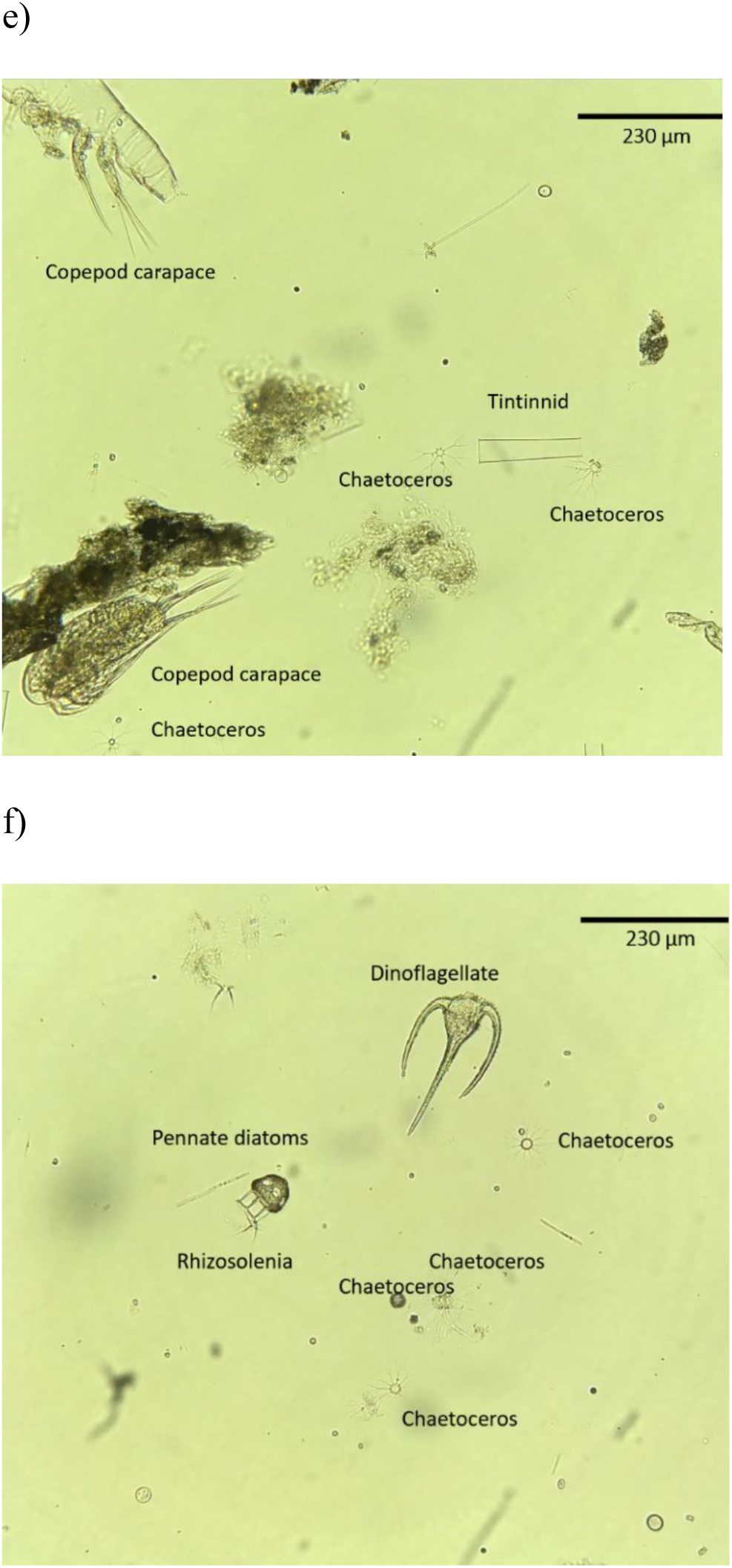
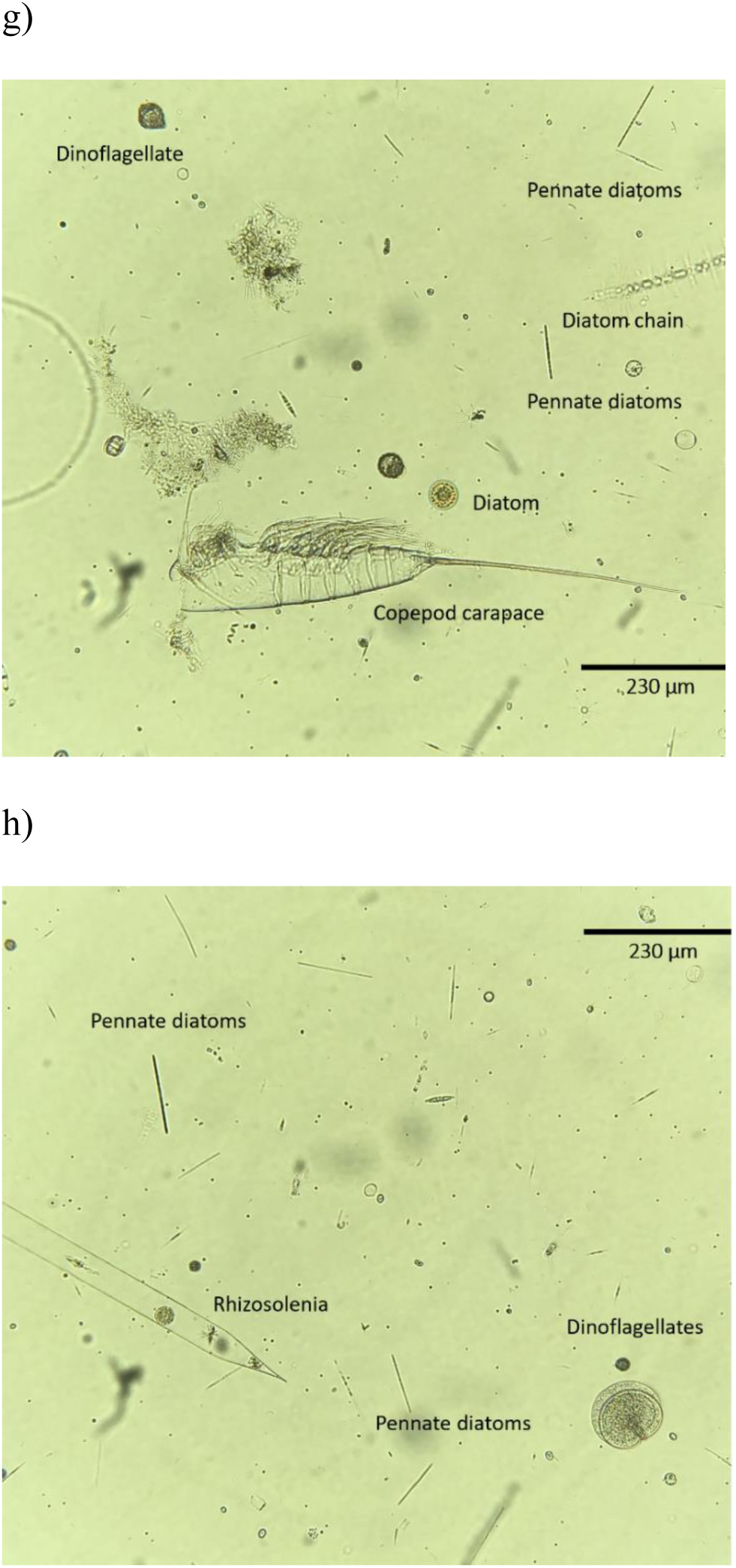
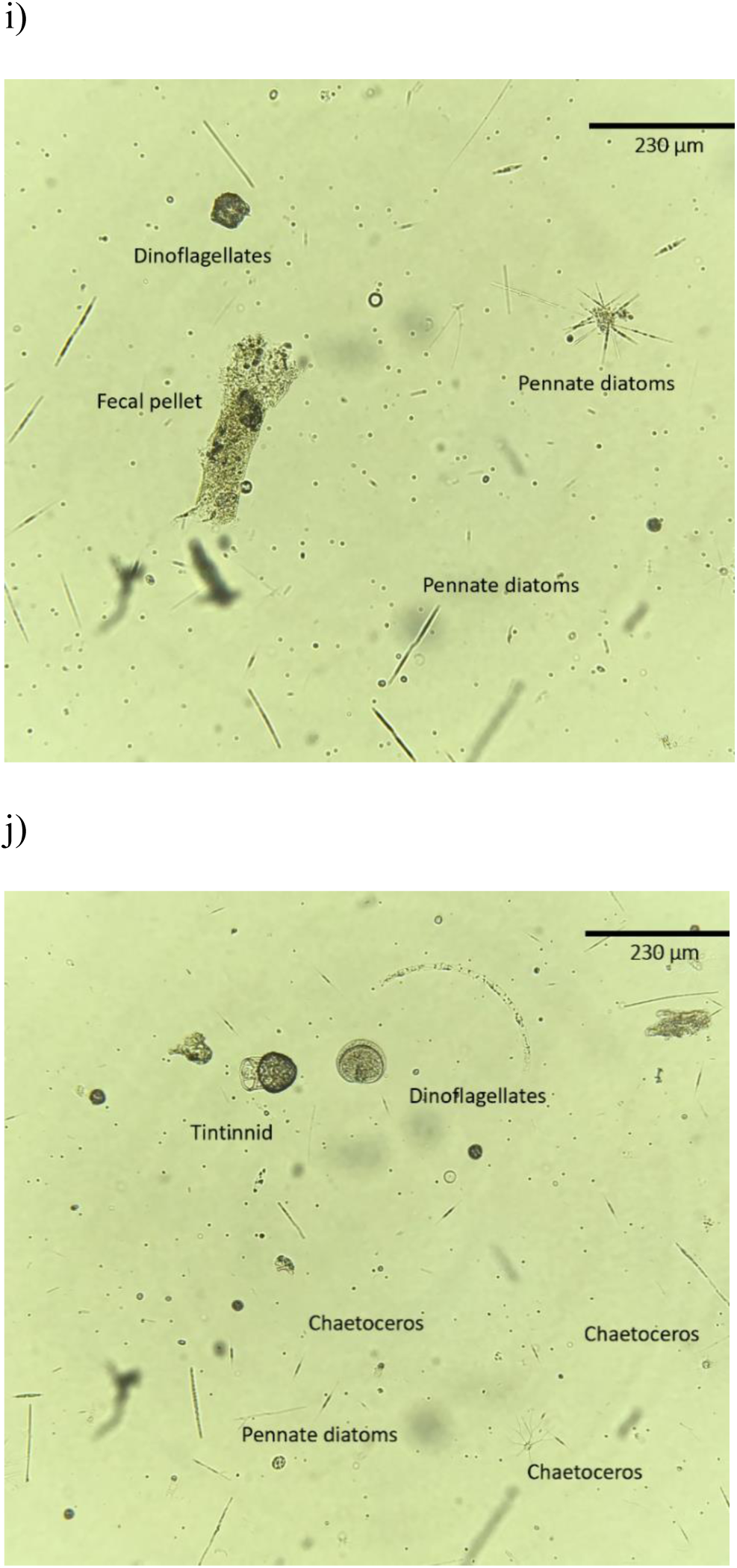
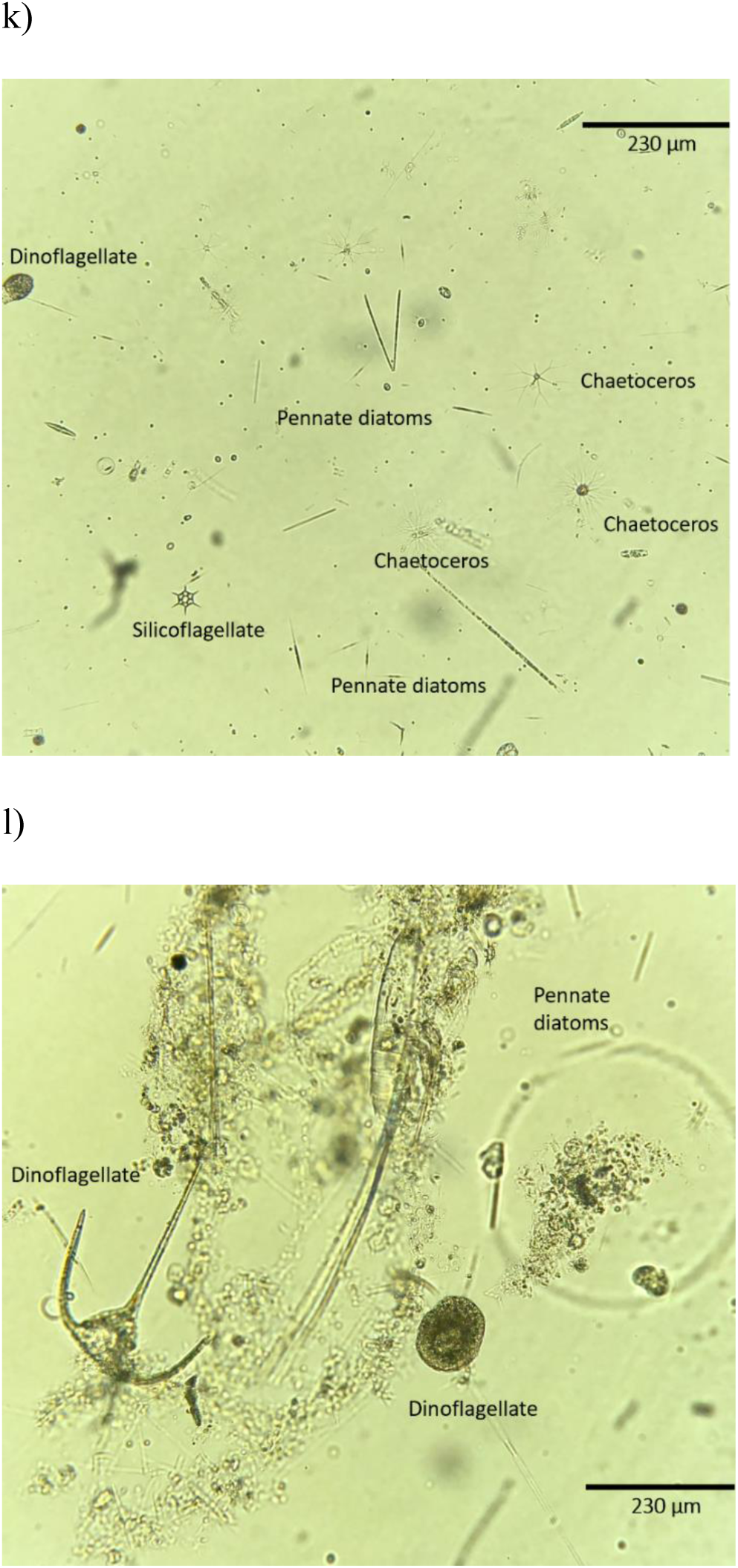
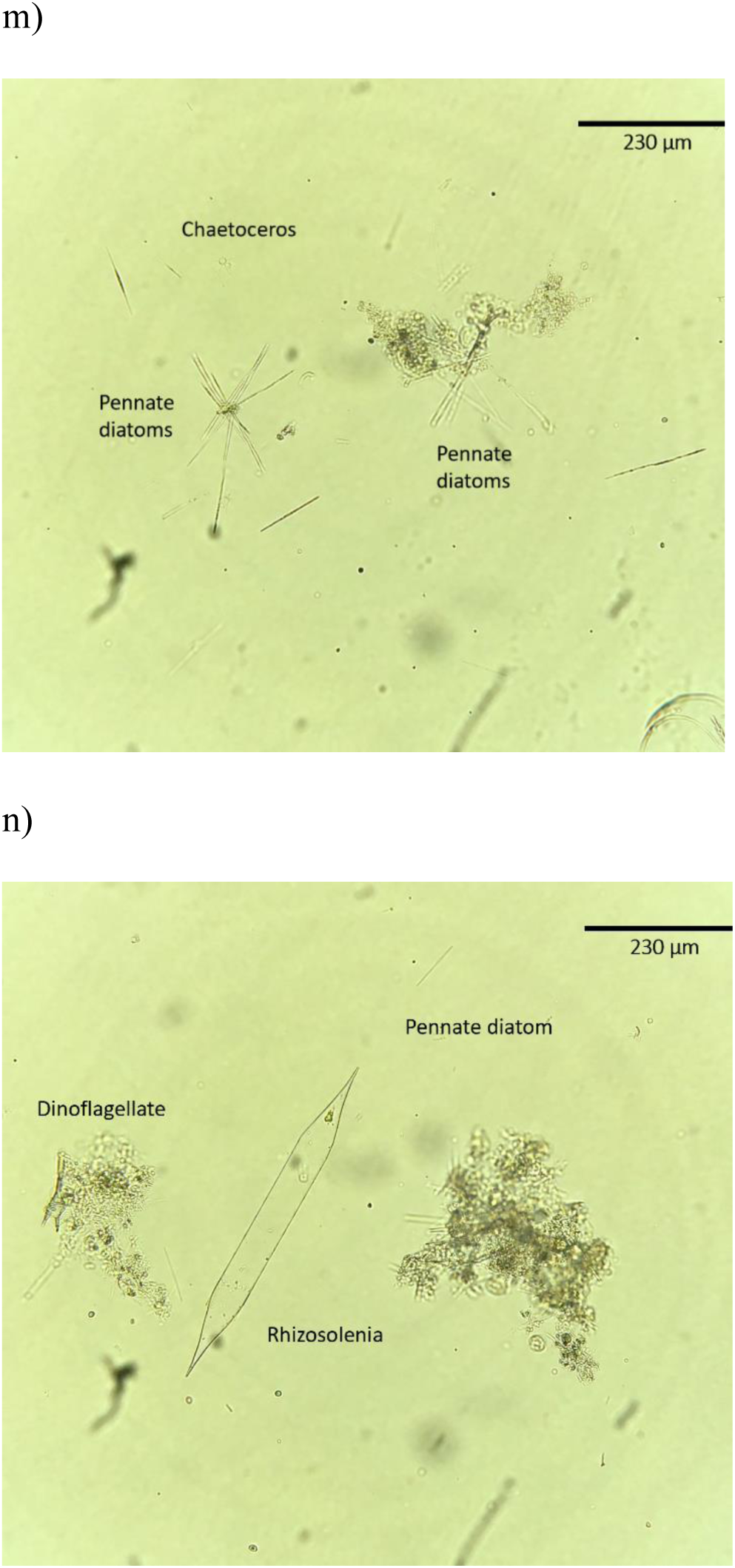
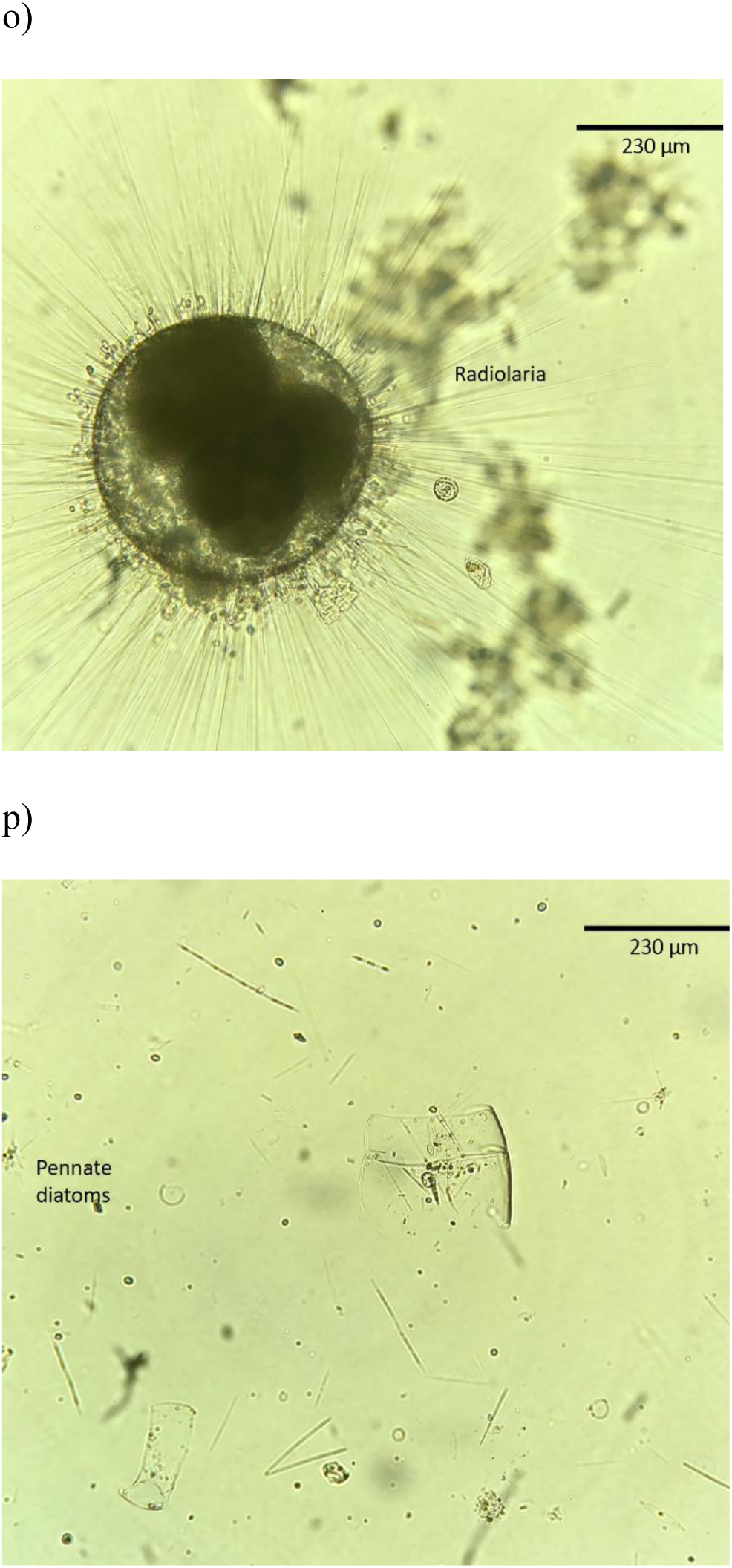
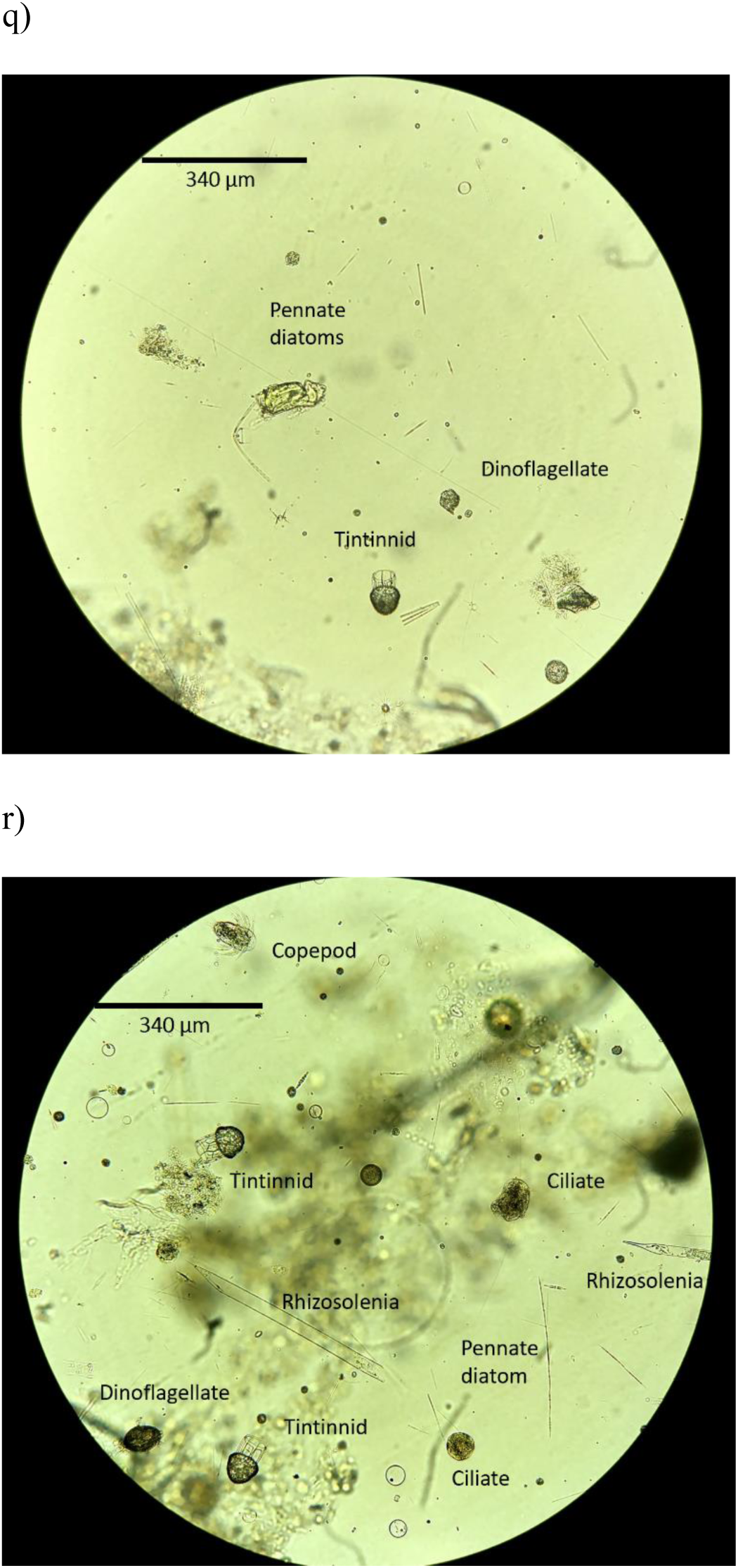
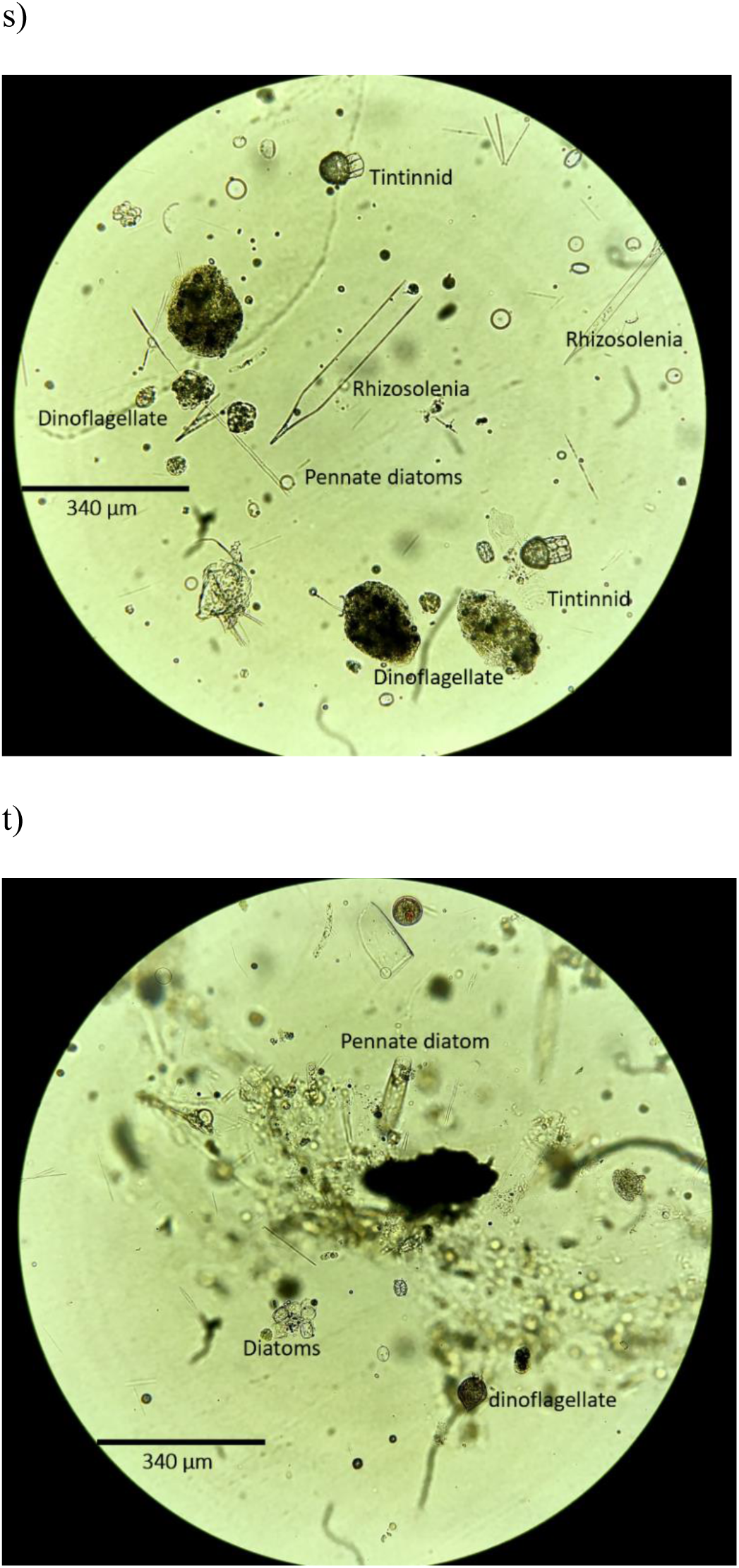
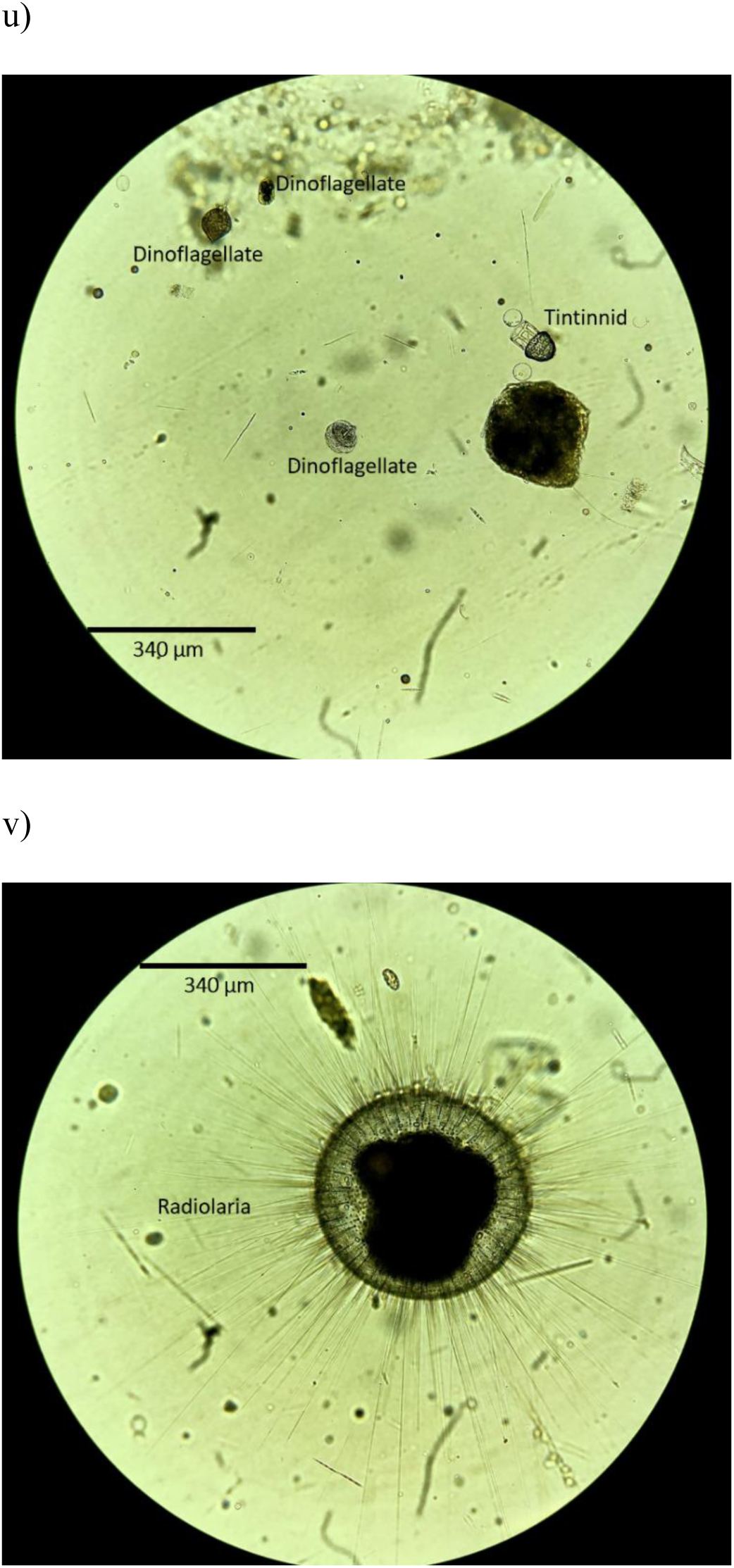
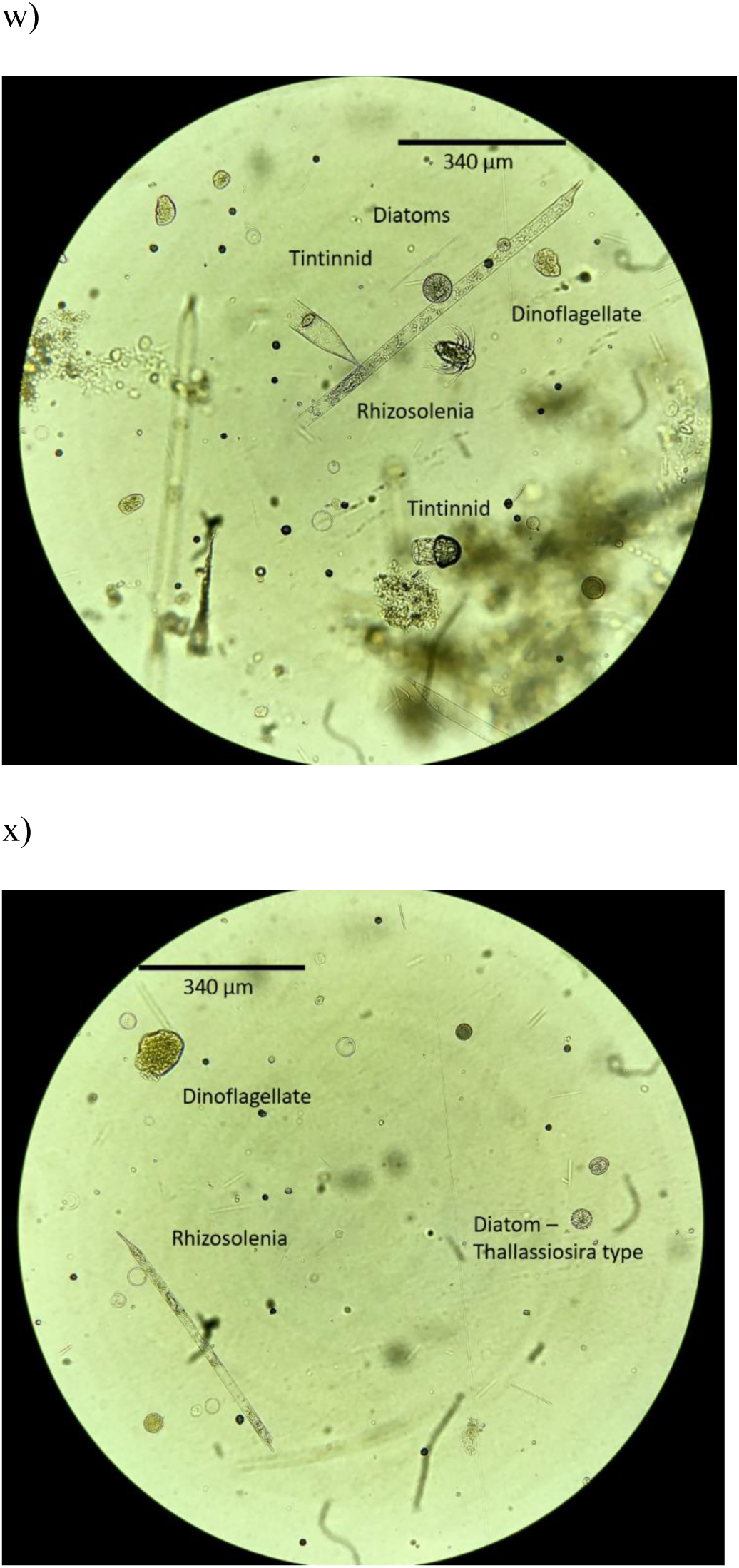
Microscopy photos with classification of plankton cells associated with marine snow collected at 50–60 m depth from the MSC trays during the field campaign on May 12 (a-f), May 17 (g-k), May 26 (l-q), and May 29 (r-x).

